# The development of intracytoplasmic membranes in alphaproteobacteria involves the conserved mitochondrial crista-developing protein Mic60

**DOI:** 10.1101/2022.06.14.496148

**Authors:** Sergio A. Muñoz-Gómez, Lawrence Rudy Cadena, Alastair T. Gardiner, Michelle M. Leger, Shaghayegh Sheikh, Louise Connell, Tomáš Bilý, Karel Kopejtka, J. Thomas Beatty, Michal Koblížek, Andrew J. Roger, Claudio H. Slamovits, Julius Lukeš, Hassan Hashimi

**Affiliations:** Ecologie Systématique Evolution, Université Paris-Saclay, AgroParisTech, Orsay, France; Institute of Parasitology, Biology Center, Czech Academy of Sciences, České Budějovice (Budweis), Czech Republic; Faculty of Science, University of South Bohemia, České Budějovice (Budweis), Czech Republic; Center Algatech, Institute of Microbiology, Czech Academy of Sciences, Třeboň, Czech Republic; Institute of Evolutionary Biology (CSIC-Universitat Pompeu Fabra), Barcelona, Catalonia, Spain; Department of Chemistry and Biomolecular Sciences, University of Ottawa, Ottawa, Canada; Department of Microbiology and Immunology, University of British Columbia, Vancouver, Canada; Centre for Comparative Genomics and Evolutionary Bioinformatics, Department of Biochemistry and Molecular Biology, Dalhousie University, Halifax, Nova Scotia, Canada

**Keywords:** *Cereibacter*, *Rhodobacter*, *Rhodopseudomonas*, chromatophores, eukaryote, endosymbosis, MICOS

## Abstract

Mitochondrial cristae expand the surface area of respiratory membranes and ultimately allow for the evolutionary scaling of respiration with cell volume across eukaryotes. The discovery of Mic60 homologs among alphaproteobacteria, the closest extant relatives of mitochondria, suggested that cristae might have evolved from bacterial intracytoplasmic membranes (ICMs). Here, we investigated the predicted structure and function of alphaproteobacterial Mic60, and a protein encoded by an adjacent gene Orf52, in two distantly related purple alphaproteobacteria, *Rhodobacter sphaeroides* and *Rhodopseudomonas palustris*. In addition, we assessed the potential physical interactors of Mic60 and Orf52 in *R. sphaeroides*. We show that the three α-helices of mitochondrial Mic60’s mitofilin domain, as well as its adjacent membrane-binding amphipathic helix, are present in alphaproteobacterial Mic60. The disruption of Mic60 and Orf52 caused photoheterotrophic growth defects, which are most severe under low light conditions, and both their disruption and overexpression led to enlarged ICMs in both studied alphaproteobacteria. We also found that alphaproteobacterial Mic60 physically interacts with BamA, the homolog of Sam50, one of the main physical interactors of eukaryotic Mic60. This interaction, responsible for making contact sites at mitochondrial envelopes, has been conserved in modern alphaproteobacteria despite more than a billion years of evolutionary divergence. Our results suggest a role for Mic60 in photosynthetic ICM development and contact site formation at alphaproteobacterial envelopes. Overall, we provide support for the hypothesis that mitochondrial cristae evolved from alphaproteobacterial ICMs, and therefore have improved our understanding of the nature of the mitochondrial ancestor.

## Introduction

Mitochondria are organelles inferred to have been present in the last common ancestor of all eukaryotes (reviewed in Roger et al. 2017). Unlike most other organelles (e.g., the endoplasmic reticulum, nucleus, cytoskeleton, etc.), the mitochondrion evolved from an endosymbiont most closely related to extant alphaproteobacteria ^2–4^. In aerobic eukaryotes, mitochondria produce most of the ATP of the cell through aerobic respiration, i.e., the harnessing of energy through the coupling of electron transport to chemiosmosis with oxygen as a terminal electron acceptor. Mitochondria also compartmentalize other metabolic pathways ^1,5^. Because aerobic respiration occurs at internalized membranes that can expand greatly, mitochondria allow for the proportional increase (or linear scaling) of respiration with cell volume across eukaryotes ^6^. Mitochondria are thus one of the innovations that likely allowed many eukaryotes to achieve larger cell volumes coupled to relatively fast growth rates, and ultimately opened new evolutionary trajectories. Elucidating the origin of mitochondria and their respiratory membranes may shed light on the origin of eukaryotic cells.

The specialization of mitochondria as respiratory organelles is most clearly reflected in their internal structure. The mitochondrial inner membrane invaginates into specialized sub-compartments called cristae, the structural hallmarks of the organelle ^7–9^. The MICOS (Mitochondrial Contact Site and Cristae Organizing System) complex and oligomers of the F_1_F_O_-ATP synthase are two of the most evolutionarily conserved factors responsible for the development and shape of cristae^10,11^. Whereas ATP synthase oligomers bend crista membranes at their rims to produce diverse crista shapes ^12,13^, the MICOS complex creates both crista junctions and contact sites that compartmentalize, stably anchor, and maintain cristae at the mitochondrial envelope ^14^. These functions of the MICOS complex appear to be largely conserved across phylogenetically disparate eukaryotes, such as in the animal *Homo sapiens*, the fungus *Saccharomyces cerevisiae*, the land plant *Arabidopsis thaliana*, and the parasitic protist *Trypanosoma brucei* ^15–18^.

Studies on the evolutionary history of MICOS revealed that this multi-protein complex is ancestrally present in all eukaryotes and predates the origin of mitochondria (Muñoz-Gómez et al. 2015; Huynen et al. 2016). The central and scaffolding subunit of the MICOS complex, the Mic60 protein, traces back to the *Alphaproteobacteria*, the group from which mitochondria descended. Indeed, Mic60 serves as a phylogenetic marker that is uniquely present in the *Alphaproteobacteria* and the mitochondrial lineage ^2^. While only the C-terminal signature mitofilin domain of Mic60 is sufficiently conserved at the sequence level, the overall predicted secondary structure of Mic60 has also been conserved in both mitochondrial and alphaproteobacterial homologs ^19^ In addition to having homologues of Mic60, many alphaproteobacteria also develop either lamellar or vesicular intracytoplasmic membranes (ICMs) that house diverse electron transport chains involved in methanotrophy, nitrification, and anoxygenic photosynthesis ^21–25^. This raises the possibility that Mic60 is involved in the development and/or stability of ICMs in alphaproteobacteria, and that mitochondrial cristae evolved from ancestral alphaproteobacterial ICMs ^26^. Support for the functional conservation of alphaproteobacterial Mic60 comes from its capacity to bind and bend membranes *in vitro* and heterologously in the gammaproteobacterium *Escherichia coli* ^27^ However, the precise role of Mic60 has not yet been directly studied in alphaproteobacteria, and thus the evolutionary relationship between cristae and ICMs remains unknown.

Though early ideas linked cristae to ICMs based simply on morphological resemblance, the iconography of the field (i.e., the aggregate of scientific diagrams) has mostly depicted cristae as post-endosymbiotic adaptations of mitochondria ^26^. To better understand the function of Mic60 in alphaproteobacteria and the origin of mitochondrial cristae, we investigated the role of *mic60*, and its adjacent gene *orf52*, in two distantly related purple alphaproteobacteria: the vesicular ICM-developing *Rhodobacter* (*Cereibacter*) *sphaeroides* (*Rhodobacterales*) and lamellar ICM-developing *Rhodopseudomonas palustris* (*Rhizobiales*). We first explored the genomic context, large-scale phylogenetic distribution, and sequence and structural conservation of alphaproteobacterial Mic60 homologues. We then experimentally investigated the effects of the disruption and overexpression of Mic60 and Orf52 in photoheterotrophic growth and ICM development. Finally, we assessed the higher-order assembly and physical interactors of alphaproteobacterial Mic60 and Orf52.

## Results

### Alphaproteobacterial *mic60* is clustered with a syntenic neighboring gene, *orf52*

A survey of the genomic context of *mic60* in several alphaproteobacterial species revealed that *mic60* is genetically linked to genes involved in the heme biosynthesis pathway. In most alphaproteobacterial genomes, *mic60* is downstream of *hemC* (hydroxymethylbilane synthase; HMBS) and *hemD* (uroporphyrinogen-III synthase; UROS), and upstream of a hypothetical protein-coding gene sometimes misannotated as *hemY*. All four genes have the same orientation and are usually tightly clustered with little intergenic space in between them, which may suggest that they are functionally related or co-transcribed as part of the same operon. This is consistent with the regulatory requirements of Mic60 according to its hypothesized function in ICM development, which requires heme biosynthesis for the proper assembly of cytochromes ^26^. Indeed, HemD is occasionally fused to Mic60 in members of the *Rhodospirilales*, *Kiloniellales*, and *Rhizobiales* (Fig. 1A) ^20^ In the yeast *S. cerevisiae*, the MICOS complex has been reported to interact with the enzyme ferrochelatase (HemH in alphaproteobacteria) which catalyzes the insertion of ferrous iron into protoporphyrin IX, the eighth and final step in heme biosynthesis ^28^.

**Figure 1.**
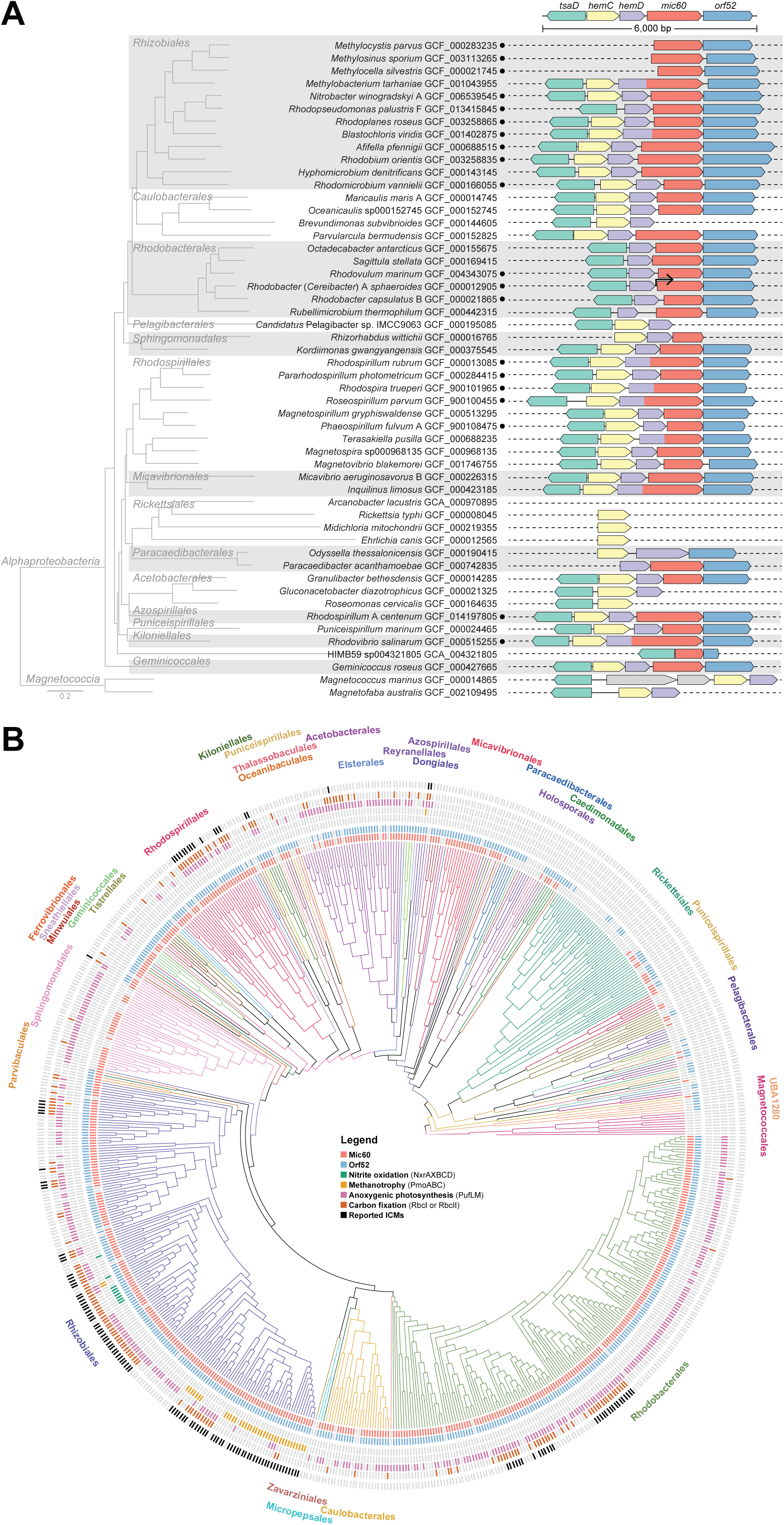
Genomic context of *mic60* and phylogenetic distribution of Mic60, Orf52, and ICMs across the *Alphaproteobacteria*. **A.** The genomic context for *mic60* in representative ICM-developing alphaproteobacteria and their relatives. Black circles to right of species identifiers denote alphaproteobacteria that develop extensive ICMs. The bent arrow denotes the TSS in *R. sphaeroides*. Representative taxa were selected manually and a supermatrix of 117 single-copy marker genes was assembled with GToTree ^34^. The compositional heterogeneity of the supermatrix was decreased with □trimmer and a phylogenetic tree was inferred with IQ-TREE and the LG+C60+F+G4 model ^35^ **B.** A comprehensive phylogenetic tree of the *Alphaproteobacteria* that displays the distributions of Mic60, Orf52, markers for ICM-associated physiologies, and reported ICMs across a maximally diverse set of taxa down sampled from the GTDB R207 database ^36^. A supermatrix of single-copy marker genes was assembled with GTDB-Tk, a phylogenetic tree was inferred with IQ-TREE (-fast mode), and the longest-branching taxa were identified and removed with TreeShrink ^37^. The down sampling was performed with Treemmer ^38^ and constrained in such a way that all taxa with ICMs and ICM-associated physiologies were retained. TreeViewer was used to display the phylogenetic distributions of traits on the tree. Protein searches were performed with hmmsearch of the HMMER suite using both Pfam and custom pHMMs ^39^.

Although *hemC* and *hemD* are genuine enzymes of the heme biosynthesis pathway, the gene downstream of *mic60* clearly does not encode a heme biosynthetic enzyme. This protein is usually misannotated as HemY because it contains a conserved hemY_N domain (PF07219) at its N-terminus. Confusingly, this domain is unrelated to genuine *hemY* (protoporphyrinogen IX oxidase), which is rather uncommon among alphaproteobacteria. Instead of *hemY*, most alphaproteobacteria use the product of *hemJ* to synthesize protoporphyrin IX (very few alphaproteobacteria have a genuine *hemY* gene homologous to that of *E. coli* and *Bacillus subtilis*) ^29^ Here, the protein encoded by the gene downstream of *mic60* will be referred to as Orf52 based on its predicted size of 52 kDa in *R. sphaeroides* 2.4.1. Like alphaproteobacterial Mic60, Orf52 is an integral membrane protein, but it possesses two transmembrane segments instead of one at its N-terminus, and seems to expose its bulk to the periplasmic space. Moreover, Orf52 contains several tetratricopeptide repeat motifs, which are usually involved in protein-protein interactions and found in proteins that are part of multiprotein complexes. In *R. sphaeroides*, the *mic60*□*orf52* gene pair is co-transcribed as indicated by a transcription start site (TSS) in the intergenic region between *hemD* and *mic60* ^30^ (Fig 2A). In the magnetosome gene island (MAI) of *Magnetospirillum gryphiswaldense, mic60* and *orf52* are also cotranscribed separately from their neighboring genes ^31^. The conserved motif order and composition of Mic60 and Orf52, as well as their transcriptional coupling, indicate that these proteins have structural roles and may physically interact with each other at alphaproteobacterial envelopes.

**Figure 2.**
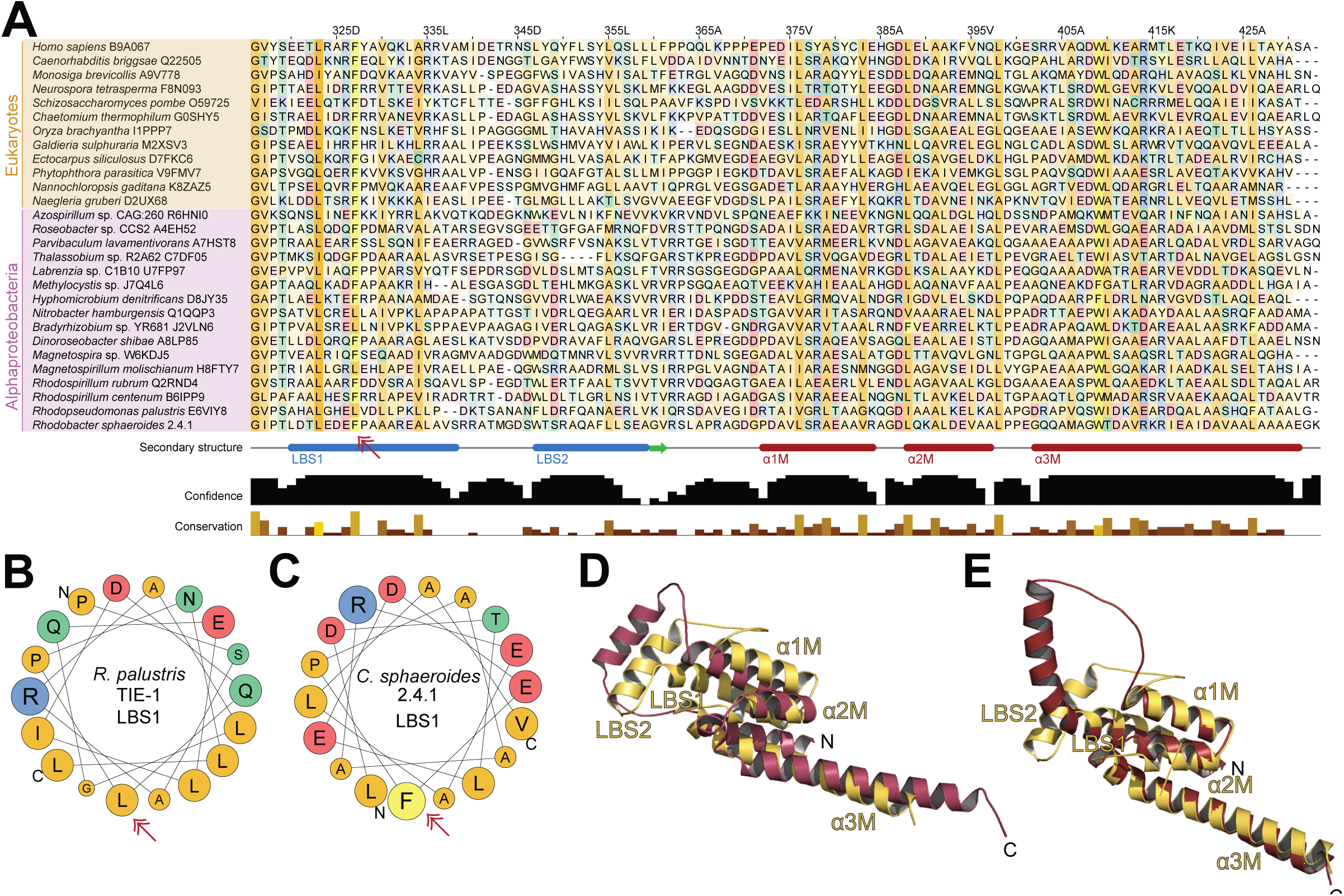
Evolutionary conservation of the secondary and tertiary structure of mitochondrial and alphaproteobacterial Mic60. **A.** Alignment of C-terminal signature mitofilin domain and its adjacent N-terminal region of representative alphaproteobacteria and eukaryotes. This alignment has the Mic60 amino acid sequence of *R. sphaeroides* as a reference and was obtained from JPred4 ^41^. The amino acid colors follow a coloring scheme based on physicochemical properties, and the intensity of the color reflect evolutionary conservation of the site in the alignment. The red arrow points to the conserved Phe327 in the *R. sphaeroides* Mic60 homolog. Common gaps are hidden from the alignment as per JPred4 output. **B.** Helical wheel projection of the amphipathic helix that comprises LBS1 in *R. palustris* as predicted by the HELIQUEST web server ^43^. **C.** Helical wheel projection of the amphipathic helix that comprises LBS1 in *R. sphaeroides* predicted as in B. **D.** Structural alignment of the C-terminal region of Mic60 homologs from the eukaryote *L. thermotolerans* (yellow) and the bacterium *R. palustris* (pink). **E.** Structural alignment of the C-terminal region of Mic60 homologs from the eukaryote *L. thermotolerans* (yellow) and the bacterium *R. sphaeroides* (red). See Fig. S1 for AlphaFold2 structure predictions of the entire protein sequences.

### Mic60 and Orf52 have broad phylogenetic distributions that overlap with the distribution of ICMs in the *Alphaproteobacteria*

Prokaryotic homologs of Mic60 have been previously shown to be restricted to alphaproteobacterial species, thus serving as a synapomorphy for the clade that comprises both mitochondria and the *Alphaproteobacteria* ^19,20,2^. Moreover, phylogenetically diverse alphaproteobacteria are known to develop extensive bioenergetic ICMs that house electron transport chains associated with physiological processes such as methanotrophy (methane oxidation), nitrification (nitrite oxidation), and (aerobic or anaerobic) anoxygenic photosynthesis ^21–23,26,24^. However, it is unclear to what extent the presence of Mic60, which has been hypothesized to be involved in ICM development ^26^, correlates with the occurrence of ICMs. To better understand the evolutionary relationship between Mic60 and ICMs, we investigated their large-scale phylogenetic distributions in the *Alphaproteobacteria*. To do this, we searched the GTDB R207 database that currently comprises more than 7,684 alphaproteobacterial genomes, with profile Hidden Markov Models (pHMMs) for Mic60, Orf52, and markers for ICM-associated physiologies (Fig. 1B).

These analyses show that both Mic60 and Orf52 have a broad and dense phylogenetic distribution that encompasses the much more sporadic distribution of reported ICMs in the *Alphaproteobacteria* (Fig. 1B). Methanotrophy and nitrite oxidation are restricted to a very few genera (e.g., *Methylocella, Methylosinus*, and *Methylocystis*, and *Nitrobacter)*, whereas photoautotrophy (or the capacity to harvest light’s energy to fix carbon dioxide) is phylogenetically widespread (Fig. 1B). The presence of these ICM-associated physiologies largely overlaps with those species reported to develop extensive ICMs (Fig. 1B). Furthermore, the prediction of a photoautotrophic physiology in several species not yet studied ultrastructurally suggests that these may also develop extensive ICMs. On the other hand, phototrophs (represented by aerobic anoxygenic photoheterotrophs) have a much broader phylogenetic distribution ^32^ (Fig. 1B), though they are not often associated with the presence of extensive ICMs. It is known, however, that aerobic anoxygenic phototrophs can develop less conspicuous ICMs under some environmental conditions ^33^ Altogether, these phylogenetic patterns suggests that both Mic60 and Orf52 are ancestrally present in the *Alphaproteobacteria* and are required by extant species that either have or lack the capacity to develop ICMs.

### The predicted secondary and tertiary structure of alphaproteobacterial Mic60 is conserved

Previous studies have shown that the overall predicted secondary structure of Mic60 is conserved both in eukaryotes and alphaproteobacteria ^19,20^. In the former, Mic60 consists of an N-terminal pre-sequence (a targeting signal to the mitochondrion), followed by a transmembrane segment, a central region with coiled-coils, and a C-terminal signature mitofilin domain. The same motifs and domains are found in the same order in the alphaproteobacterial Mic60 homolog, except for the N-terminal pre-sequence which is missing, as expected. Recently, mitochondrial Mic60 has been shown to deform membranes, and thus likely to introduce curvature at crista junctions ^40,27^. This membrane-deforming capability depends on a lipid-binding site (LBS) that is found in between the central coiled-coils and the C-terminal mitofilin domain. This LBS comprises two α-helices (LBS1 and LBS2), the first of which is amphipathic and presumably inserts itself into the mitochondrial inner membrane ^40^ LBS1 is extremely important for the function of Mic60 as its removal or mutation leads to the loss of membrane binding and deformation, and also to phenotypes quite similar to those obtained when the entire *MIC60* gene is deleted in *S. cerevisiae* ^40^.

To investigate whether the membrane-bending α-helices of eukaryotic Mic60 are present in its alphaproteobacterial homologs, we first performed pHMM-sequence searches against the UniProtKB database. We also predicted the secondary structures of both eukaryotic and alphaproteobacterial Mic60 homologs with JPred4 ^41^. A detailed inspection of both the alignments and predicted secondary structures revealed that alphaproteobacterial Mic60 has retained the two α-helices that comprise the LBS of eukaryotic Mic60 (Fig. 2A). Helical wheel projections further show that LBS1 is amphipathic in both *R. sphaeroides* and *R. palustris* (Fig. 2B, C). In addition, the functionally critical amino acid position Phe573 in the yeast *Chaetomium thermophilum* ^40^ is largely conserved among alphaproteobacteria (e.g., Phe327 in *R. sphaeroides;* see Fig. 2A).

Next, we predicted the tertiary structure of the Mic60 homologs of the yeast *Lachancea thermotolerans*, whose crystal structure was recently partially solved ^42^, and the alphaproteobacteria *R. sphaeroides* and *R. palustris* using AlphaFold2 (Fig. S1A-D). The predicted structures confirm the presence of the two α-helices (LBS) in the linker region between the central coiled-coils and the C-terminal mitofilin domain (Fig. S1A-D). Furthermore, the predicted structures show that the three conserved α-helices that comprise the mitofilin domain of eukaryotic Mic60 (α1-3M), as well as the last two small α-helices of the central coiled-coil region (α2-3C), are also present in alphaproteobacterial Mic60 (Fig. S1A-D). Structural alignments reveal that the C-terminal region of alphaproteobacterial Mic60 (i.e., LBS+mitofilin) largely overlap with that of its eukaryotic homolog (Fig. 1D, E). The major structural differences between eukaryotic and alphaproteobacterial Mic60 homologs are that the former is, on average, a longer protein with a larger segment of central coiled coils, and has a transmembrane segment much closer to the N-terminus of the protein (Fig. S1A-D). The agreement between the predicted AlphaFold2 structure with high-confidence pLDDT scores and the partially experimentally solved structure at amino acids 207-382 ^42^ suggests that eukaryotic Mic60 indeed folds into a long α1C helix (Fig. S1B). On the other hand, alphaproteobacterial Mic60 may similarly have a long α1C helix, but the lower pLDDT scores in this region of the predicted AlphaFold2 structure make it currently uncertain (Fig. S1D). In summary, the above observations indicate that alphaproteobacteria Mic60 (1) has a largely conserved secondary and tertiary structure relative to eukaryotic Mic60, and (2) contains a conserved amphipathic LBS1 helix that likely aids in membrane-binding and -bending, as demonstrated *in vitro* for its eukaryotic ortholog ^27,40^.

### Knock out of *mic60* and *orf52* significantly affects photoheterotrophic growth in *R. sphaeroides* and *R. palustris*

Purple alphaproteobacteria develop extensive intracellular membranes (ICM) in the presence of light and the absence of oxygen ^21,44,22^. These ICMs house the photosynthetic apparatus and electron transport chain, which is generally composed of light-harvesting complexes 1 and 2 (LH1 and LH2), a type II reaction center (RC), a cytochrome *bc*_1_, a periplasmic cytochrome *c*_2_, and an ATP synthase ^45^. By means of cytochrome *bc*_1_, which is also shared with the respiratory chain, the photosynthetic chain creates a proton motive force across the ICM that is harvested by the ATP synthase to produce ATP ^22^. ICMs are often continuous with, but sometimes detached from ^46^, the cytoplasmic membrane (CM) ^47^, just as cristae are continuous with the mitochondrial inner membrane. In *S. cerevisiae* mitochondria, the disruption of Mic60 leads to functional defects such as decreased growth rate under respiratory conditions (i.e., non-fermentable media) and increased production of oxygen radicals ^48,49^. If Mic60 is involved in the development of photosynthetic ICMs, then its disruption should lead to growth defects in the absence of oxygen and the presence of light (i.e., photosynthetic conditions).

To assess whether Mic60 and Orf52 have an impact on photoheterotrophic growth and ICM development, we first knocked out these genes in two phylogenetically distant purple alphaproteobacteria amenable to reverse genetics, *R. sphaeroides* and *R. palustris* ^50,51^. To create knockout strains, suicide plasmids containing the knockout gene construct flanked by homologous stretches were transferred into both species via conjugation using a suitable *E. coli* strain. After selection and counter-selection for a first and second recombination events (to respectively insert and excise the suicide plasmid into the host genome), these knockout strains *(Δmic60* and *Δorf52)* were confirmed by PCR assays using several sets of internal and external primers that flank the knockout gene construct junctions (Fig. S2A-B).

Under chemoheterotrophic conditions (i.e., absence of light and presence of oxygen), purple alphaproteobacteria do not develop photosynthetic ICMs for light harvesting. Our data show that there are no growth differences between *mic60^+^orf52^+^* wild type (WT) and the Δ*mic60* and Δ*orf52* strains under these conditions (Fig. S3A). On the other hand, these bacteria develop a moderate amount of ICMs under anoxia and high light intensity. The Δ*mic60* strains had a slower photoheterotrophic growth rate relative to WT in both *R. sphaeroides* and *R. palustris* (Fig. 3A, 3C). Under these conditions, the Δ*orf52* decreased photoheterotrophic growth significantly in *R. palustris* but not in *R. sphaeroides*. Under anoxia and low light intensity, purple alphaproteobacteria upregulate the LH2 complex and develop even larger amounts of ICMs to increase light capturing. As expected, both Δ*mic60* and Δ*orf52* strains displayed even slower photoheterotrophic growth rates in both *R. sphaeroides* and *R. palustris* in low light, when ICM development increases (Fig. 3B, 3D. Interestingly, Δ*orf52* displayed much more severe decreases in growth rate, and even lower yield at stationary phase, than Δ*mic60* in *R. palustris* (Fig. 3B). This is opposite from what was observed in *R. sphaeroides* (Fig. 3D) and indicates that these proteins might contribute differently to photoheterotrophic growth in these two distantly related species. In summary, the slower photoheterotrophic growth rate of the knockout strains, especially under low light when larger amounts of ICMs are required, suggests that both Mic60 and Orf52 affect the development of photosynthetic ICMs.

**Figure 3.**
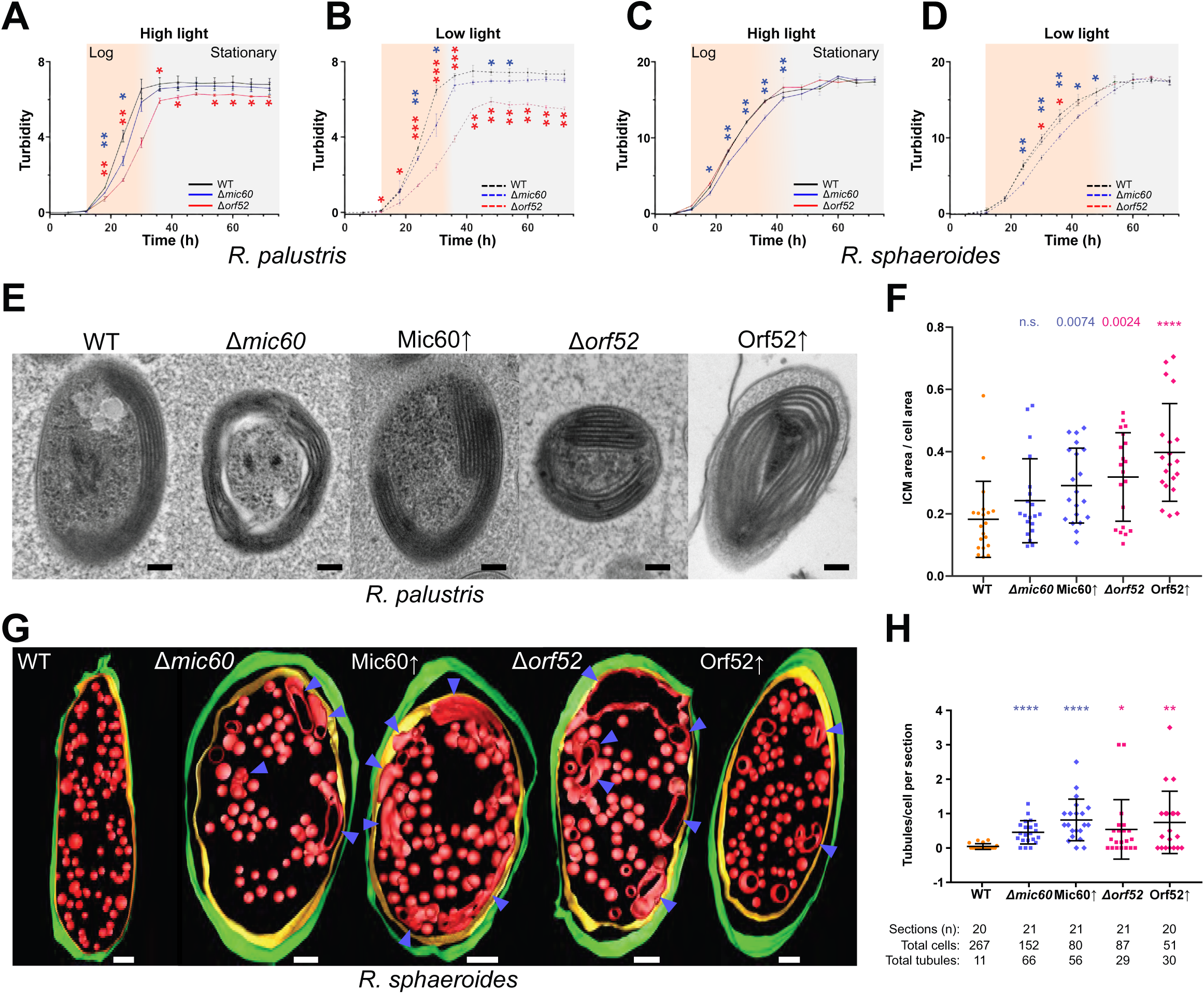
The disruption of the *mic60* and *orf52* genes causes defects in photoheterotrophic growth and) ICM development. **A-D.** Growth curves of *R. palustris* WT and Δ*mic60* and Δ*orf52* strains under high (**A**) and low (**B**) light and *R. sphaeroides* under high (**C**) and low (**D**) light. Turbidity was measured at 565 nm and expressed in arbitrary units given on the *y*-axis. Time points are expressed in hours on the *x*-axis. Growth stages are color-coded according to the labels at the top. Colored asterisks show statistical significance of differences at each time point between WT and either Δ*mic60* (blue) or Δ*orf52* (red). **E.** Exemplar TEM micrographs of each *R. palustris* strain grown in low light. **F.** Column scatter plot of the ratio of *R. palustris* ICM area/whole cell area (*y*-axis) measured from 20 cells imaged using TEM in *R. palustris* WT, knockout (Δ*mic60* and Δ*orf52)* and overexpression (Mic60↟ and Orf52↟) strains. Mean and standard deviation shown by middle bar and whiskers, respectively. Values and asterisks above each column represent statistical significance of the difference of each population in comparison to WT. **G.** Exemplar electron tomograms of each *R. sphaeroides* strain grown in low light. Blue arrows point at aberrant ICMs. **H.** Column scatter plot of tubules/cell per section (*y*-axis) for *R. sphaeroides* WT, knockout (Δ*mic60* and Δ*orf52)* and overexpression (Mic60↑ and Orf52↑) strains. The number of sections, cells and tubules for each cell lines are given in the table below the *x*-axis. Mean and standard deviation shown by middle bar and whiskers, respectively. Asterisks above each column represent statistical significance of the difference of each mutant population in comparison to WT. Related to Movie S1-S4. Scale bars in (**E)** and (**G)**, 100 nm. Statistical significance: *, P<0.05; **, P<0.01; ***, P<0.001; ****, P<0.0001. See also Fig. S1-3.

### Mic60 and Orf52 are involved in the formation of lamellar and vesicular ICMs

The disruption of mitochondrial Mic60 leads to structural defects such as the loss of crista junctions, the detachment of cristae from the mitochondrial envelope, and elongated crista membranes ^52,18^ Hypotheses about the role of Mic60 in ICM development postulate that this protein might be responsible for creating ICM junctions and contact sites that respectively compartmentalize ICMs and anchor them to the bacterial envelope ^26^. The loss and overexpression of Mic60 and Orf52 from ICM-developing alphaproteobacteria might thus lead to morphological defects in ICMs.

To directly address whether the gene products of *mic60* and *orf52* are involved in ICM formation, we generated *R. palustris* and *R. sphaeroides* strains capable of overexpressing Mic60 and Orf52 (Mic60↑ and Orf52↑, respectively). These strains were verified by reverse transcription quantitative PCR (RT-qPCR) upon induction for six hours with 1 mM isopropyl-β-D-thiogalactoside (IPTG). Both Mic60↑ and Orf52↑ strains had an increase in gene expression of ~7□9-fold in *R. sphaeroides*, and of about two-fold in *R. palustris*, relative to ex-conjugants grown in the absence of IPTG (Fig. S1C). The knockout and WT strains were grown under photoheterotrophic conditions and low light, and were IPTG-induced for six hours (in case of overexpression ex-conjugants) to maximize ICM development. After harvesting, the strains were cryopreserved, contrasted, and imaged using transmission electron microscopy (TEM). Randomized TEM micrographs were blindly scored to quantitatively evaluate phenotypes associated to the peripheral stacked lamellar ICMs of *R. palustris* and the uniformly distributed vesicular ICMs of *R. sphaeroides* ^24^ (Fig. 3E–H, Movie S1-4).

In *R. palustris*, the area occupied by the lamellar ICMs increased relative to the whole cell area in both knockout (Δ*mic60* and Δ*orf52)* and overexpression (Mic60↑ and Orf52↑) strains, as compared to the WT (ratio ICM area/cell area=0.18±0.03) (Fig. 3E–F). This increase was statistically significant in Mic60↑ (0.29±0.03) but not in Δ*mic60* (0.24±0.03). Both Δ*orf52* (0.32±0.03) and Orf52↑ (0.40±0.04) demonstrated higher and statistically significant increases in ICM:cell area ratios when compared to their Mic60 counterparts. In *R. sphaeroides*, we scored for the appearance of elongated ICMs (i.e., tubules) per cell as these were rarely observed in the WT strain (tubules/cell/section=0.04±0.02) (Fig 3G–H). Both Δ*mic60* (0.46±0.07) and Mic60↑ (0.82±0.13) showed a statistically highly significant increase in the number of tubules, whereas these structures were observed to a lesser extent in Δ*orf52* (0.54±0.19) and Orf52↑ (0.74±0.20); the most conspicuous outliers (≥3 tubules/cell/section) were observed in the latter cells. Moreover, among tubulated ICMs, Mic60↑ displayed a significantly higher incidence of branching ICMs relative to Δ*mic60* (Fig. S4), reminiscent of the branched cristae seen in *S. cerevisiae MIC60*↑ ^52^. Electron tomograms were used to render representative images for *R. sphaeroides* cells whose ICM membranes were not well contrasted (Fig 3G, Movie S1-4). This revealed the presence of elongated ICMs of various lengths and volumes in all the mutant strains assayed.

The ultrastructural defects of the Δ*mic60* and Δ*orf52* strains of *R. sphaeroides* and *R. palustris* are consistent with their lower photoheterotrophic growth dynamics. Namely, the Δ*orf52* strain exhibits more pronounced phenotypes than the Δ*mic60* strains in *R. palustris*, whereas the opposite is true in *R. sphaeroides*. These defects in ICM area and shape were not paralleled by the absorbance spectra of whole-cell protein extracts in *R. palustris* which suggests that the regulation of RC-LH1 and LH2 complexes is largely unaffected (Fig. S3B-C). We also observed that both the knockout and overexpression strains led to ICM area expansion. It is possible that both the disruption and overproduction of Mic60 lead to unregulated membrane growth that, under the image analyses employed here, produce seemingly similar phenotypes. However, improved sample preservation and larger-scale volumetric electron microscopy will be required to quantitatively assess changes in the shape and number of ICM junctions, which potentially differ between knockout and overexpression strains. The expansion of photosynthetic ICMs is similar to the enlarged respiratory cristae observed in *S. cerevisiae* and *T. brucei* mitochondria defective of Mic60, and to the enlarged magnetosomes reported in the magnetotactic alphaproteobacterium *M. gryphiswaldense* upon deletion of the *mic60* paralog found within the magnetosome gene island (see Discussion). Together, the ultrastructural defects displayed by both Δ*mic60* and Δ*orf52* suggest that these genes are involved in the development of photosynthetic ICMs in purple alphaproteobacteria.

### Mic60 and Orf52 are assembled into a 250 kDa complex in the ICMs of *R. sphaeroides*

Eukaryotic MICOS comprises 6-9 subunits in *H. sapiens, S. cerevisiae*, and *T. brucei*. Four of them, namely Mic60, Mic10, Mic19 and Mic12, are inferred to have been ancestral to eukaryotes ^19,20^. Furthermore, the MICOS complex has been suggested to engage in interactions with a myriad of other proteins, such as the outer membrane β-barrel insertase Sam50, the outer membrane translocase Tom40, and the mitochondrial intermembrane space assembly protein Mia40 ^13,53^. It is thus possible that alphaproteobacterial Mic60 and Orf52 are part of a larger multi-protein complex at alphaproteobacterial envelopes.

To investigate whether Mic60 and Orf52 form part of a multi-protein complex, *R. sphaeroides* was conjugated with a suitable *E. coli* strain for the transfer of plasmids that allow the IPTG-inducible expression of either Mic60 or Orf52 bearing a C-terminal hexa-histidine (6xHis) tag. The expression of Mic60-6xHis and Orf52-6xHis was verified by Western blot analysis using anti-6xHis antibodies (Fig. 4A). Mic60-6xHis was found to migrate as a ~62 kDa protein, in contrast to its theoretical 43.8 kDa molecular weight; this is likely explained by the anomalous mobility of acidic proteins in SDS-PAGE ^54^. The two overexpression strains, i.e., Mic60-6xHis↑ and Orf52-6xHis↑, alongside the Mic60^+^Orf52^+^ (WT) control strain, were disrupted by high pressure homogenization to allow for ICM isolation (i.e., chromatophores) by differential centrifugation. The absorption spectra of the isolated ICMs from each strain were essentially identical, which indicates that the overexpression of Mic60-6xHis and Orf52-6xHis does not affect the RC-LH1:LH2 protein composition ratio of ICMs (Fig. 4B).

**Figure 4.**
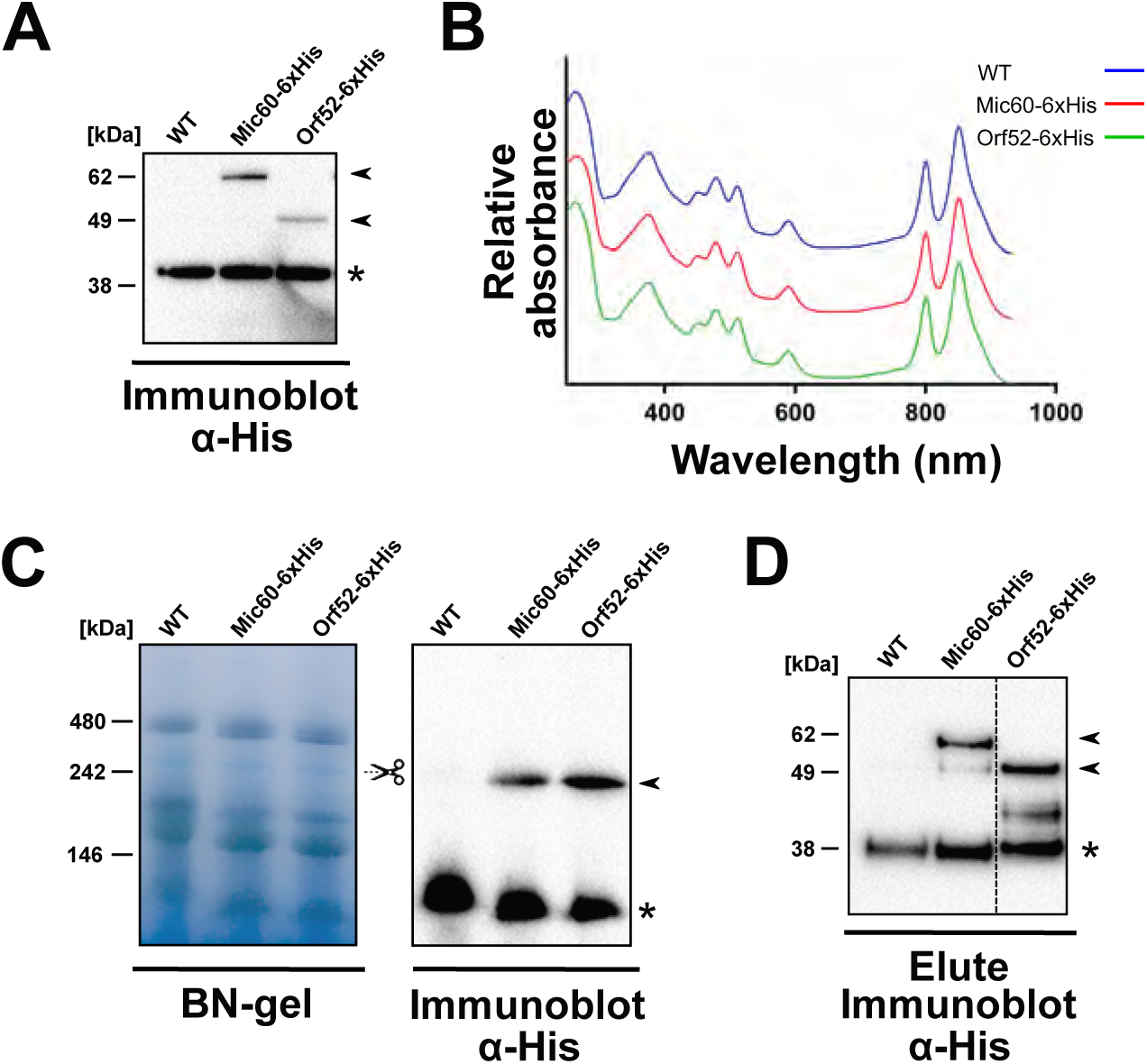
Mic60 and Orf52 assemble into a 250 kDa protein complex in *R. sphaeroides*. **A.** Immunoblot verifying the expression of Mic60-6xHis and Orf52-6xHis. **B.** Absorption spectra (~280-950 nm scan, *x*-axis) of ICMs isolated from Mic60-6xHis, Orf52-6xHis, and WT control strains. Relative absorbance given on *y*-axis and spectra are normalized to the bacteriochlorophyll *a* Qx peak at 590 nm. Spectra are stacked on top of each other and color-coded according to the legend on the upper right-hand corner. **C.** Protein complexes from detergent-solubilized isolated ICMs as resolved by BN-PAGE. Scissor symbol indicates the ~250 kDa band excised for MS analysis on left gel (see also Fig. S3A and Dataset 1). Right gel shows immunoblot demonstrating that Mic60-6xHis and Orf52-6xHis assemble into a ~250 kDa band. **D.** Immunoblot of Mic60-6xHis and Orf52-6xHis AP eluates showing that the 6xHis-tagged bait proteins were successfully purified from detergent-solubilized isolated ICMs. For **A**, **C**, and **D**, molecular weight markers shown on left. All immunoblots use arrows to point at specific antibody signals from Mic60-6xHis and Orf52-6xHis (absent for WT) controls whereas asterisks denoted non-specific band used as loading control.

The isolated ICMs were resolved in blue native PAGE (BN-PAGE) gels and transferred onto a membrane for probing with an anti-His antibody (Fig. 4C). Both Mic60-6xHis and Orf52-6xHis incorporate into a ~250 kDa multi-protein complex, with the antibody signal absent from the WT lane. Notably, the immunoreactive ~250 kDa-sized band seems to correspond to a faint and similarly sized Coomassie-stained band in the BN-PAGE gel. To identify proteins migrating in this region of the BN-PAGE gel, four ~250 kDa bands were excised from *R. sphaeroides* WT and the proteins eluted and analyzed by liquid chromatography-tandem mass spectroscopy (LC-MS/MS) (Fig. S2A). Mic60 and Orf52 were found among the 175 top proteins in the ~250 kDa band with a mean intensity score >23 in all four independent biological replicates (Dataset 1). The top-three hits corresponded to the RC complex subunits H, M, and L, which were previously shown to co-migrate with Mic60 in the BN-PAGE gels ^55^ These data are consistent with both Mic60 and Orf52 previously being detected in isolated ICMs and/or in their developmental precursors, i.e., ‘upper pigmented bands’ (UPB) ^55,56^. These experiments thus show that both Mic60 and Orf52 are part of a higher-order assembly complex of ~250 kDa present in photosynthetic ICMs.

### Mic60 and Orf52 interact with the outer membrane β-barrel insertase BamA

Among several interactions reported for the eukaryotic MICOS complex (e.g., with Tom40, Sam50, Mia40), it was suggested that the Mic60-Sam50 interaction may have predated the origin of mitochondria ^53^. This interaction is required for making contact sites at the mitochondrial envelope of phylogenetically disparate eukaryotes such as *H. sapiens, S. cerevisiae*, and *T. brucei* ^15,16,18^, and the bacterial homolog of Sam50, BamA, is a ubiquitous protein of bacteria surrounded by two membranes (diderms). In these bacteria (e.g., *Proteobacteria)*, BamA is required for the assembly of β-barrels in the outer membrane ^57–59^. Furthermore, *mic60* is genetically linked to *orf52* and co-transcribed as part of the same operon ^30^. These observations and inferences prompted us to next investigate potential physical interactors of alphaproteobacterial Mic60 (YP_351551.1) and Orf52 (YP_351550.1).

We isolated ICMs from *R. sphaeroides* and performed affinity purifications (AP) of Mic60-6xHis and Orf52-6xHis. The ICMs were solubilized with a mixture of digitonin (1.5% w/v) and Triton X-100 (0.1% v/v), and the cleared lysate was incubated with cobalt-coated Dynabeads to capture the 6xHis-tagged baits. After extensive washing, the successful capture of Mic60-6xHis and Orf52-6xHis was confirmed on a portion of beads taken for Western blot analysis with an anti-His antibody (Fig. 4D)—a mock AP was done on *R. sphaeroides* WT as a control for non-specific binding. The remaining beads from triplicate APs of Mic60-6xHis, Orf52-6xHis, and the mock controls were trypsinized and analyzed by LC-MS/MS. Protein enrichment in bait-APs in comparison to the mock control was quantified using label-free quantification as previously described ^60^. In each AP, 497 high-confidence proteins were found (Dataset 2) based on mean Andromeda confidence scores >100 in three biological replicates ^61^. Among these, proteins were considered to have true physical interaction if they (1) had Log_2_-transformed fold-enrichment and p-values larger than three (Fig. 4A–B), (2) were part of the higher-order ~250 kDa complex (see above; Dataset 1), (3) were previously reported in ICM and/or upper pigmented bands (or UPBs, which are ICM precursors; Fig. 5C–D), and (4) were predicted to have transmembrane segments or a β-barrel structure (Fig. 5C–D).

**Figure 5.**
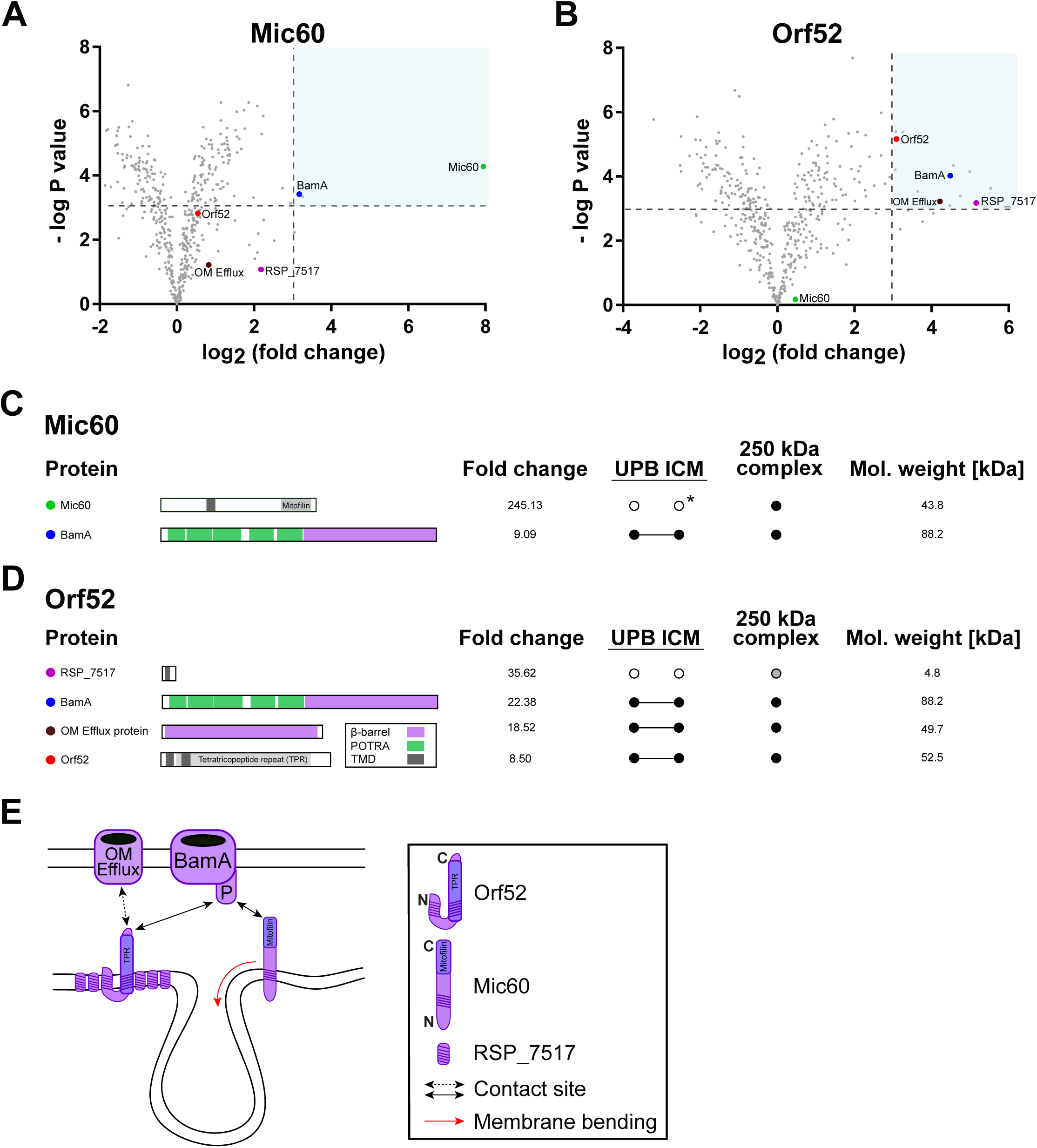
The interactomes of Mic60 and Orf52 demonstrate that both proteins interact with the outer membrane β-barrel protein BamA in *R. sphaeroides*. **A-B.** Volcano plots of proteins identified by MS in AP eluates of Mic60-6xHis (**A**) and Orf52-6xHis (**B**), with -Log P-values for each protein (*y*-axis) plotted against Log_2_-tranformed fold-enrichment over WT negative control (*x*-axis); thresholds for each value for a protein considered enriched interactors are indicated by dotted lines, demarking the shaded area of each plot. Baits and enriched interactors are indicated by labelled and colored large dots, whereas other proteins are demarked by small, grey dots. **C-D.** Summary of enriched interactors found in Mic60-6xHis (**C**) and Orf52-6xHis (**D**) APs showing the same color scheme as in **A** and **B**. From the left of protein names: schemas depicting the domain architecture of each interacting protein; its fold change in APs compared to WT negative controls; presence in upper pigmented band (UPB) and ICM proteomes (Jackson et al., 2012); presence in ~250 kDa band proteome (Dataset 1); theoretical molecular (Mol.) weight. Black dots indicate presence in the proteome; grey dot indicates presence below our defined threshold; *, presence in ICM according to MS analysis of D’Amici *et al*. (2010). **E.** Schema of Mic60 and Orf52 interactions in *R. sphaeroides*. Box contains a key with red arrow indicating an already *in vitro* membrane-remodeling activity ^27^ Related to Fig. S2 and Dataset 2.

The data show that both Mic60-6xHis and Orf52-6xHis interact with BamA (Fig. 5, S2), the β-barrel insertase subunit of the β-barrel assembly machine (BAM) complex ^59^. Somewhat unexpectedly, Mic60-6xHis and Orf52-6xHis do not interact with each other, as neither protein was found to co-purify with either bait protein (Fig. 5A–B). We cannot rule out that the C-terminal 6x-His tag may have interfered with their interaction, although this is less likely due to its small size. Mic60 was also not found to interact with the enzyme HemD, although these proteins have fused in some alphaproteobacteria (Fig 2A) ^20^. This was not unexpected as the HemD-Mic60 fusion protein is predicted to be membrane-anchored at its midpoint, having the HemD fragment exposed to the cytoplasm and Mic60 fragment on the periplasm. Moreover, the high degree of enrichment of Mic60 relative to BamA (Fig 5B, 5D) is consistent with the idea that this protein engages in homotypic interactions and forms homooligomers as its eukaryotic counterparts ^40,48,42^

In contrast to Mic60, which exhibits a strong interaction solely with BamA, Orf52 also interacts with a β-barrel protein annotated as Outer Membrane Efflux (OM Efflux) and a very small 4.8 kDa protein with an N-terminal TMD that is encoded by the RSP_7517 locus (Fig. 5A, 5C; Fig. S3D). The former belongs to the TolC protein family and was found to be more abundant in ICM than the UBP precursor ^56^, whereas the latter is among the most enriched proteins in the Orf52 AP. The enrichment of RSP_7515 is surprising as it was not previously detected in ICMs or UPBs (Jackson et al., 2012). It was, however, detected in the ~250 kDa BN-PAGE band, albeit below our defined threshold by a single peptide (Dataset 1). This may be explained by the fact that this protein is highly hydrophobic and would only yield two peptides by trypsinization, making it less amenable to detection by shotgun LC-MS/MS as employed above for the 250 kDa complex ^62^ (Fig. S3D). In conclusion, the Sam50 homolog BamA appears to be the only physical interactor of alphaproteobacterial Mic60 (Fig. 5E), and this interaction has been conserved despite ~2 billion years of evolutionary divergence of modern alphaproteobacteria from mitochondria.

## Discussion

The overarching aim of this study was to investigate the structure and function of alphaproteobacterial Mic60, thereby testing the hypothesis that ICMs were transformed into cristae during the early evolution of mitochondria ^26^. To this end, we focused on two distantly related purple alphaproteobacteria, *R. palustris* and *R. sphaeroides*. We showed that alphaproteobacterial Mic60 (1) has a conserved predicted tertiary structure and membrane-bending amphipathic helix, (2) affects photoheterotrophic growth and ICM development and shape, (3) is part of a higher-order 250 kDa multi-protein complex in photosynthetic ICMs, and (4) most likely physically interacts with the core BamA subunit of the outer membrane BAM complex. Furthermore, we also showed that Orf52 (whose gene is adjacent to in alphaproteobacteria but has no homologs in eukaryotes), also affects photoheterotrophic growth, ICM development and shape, and, although it does not interact with Mic60, also physically interacts with BamA.

These findings are consistent with previous observations and experiments that also support a functional conservation of Mic60 and its involvement in the development of both vesicular and lamellar ICMs. First, the expression profile of Mic60 follows the development of photosynthetic ICMs in the absence of oxygen and presence of light ^63^. Second, Mic60 localizes to photosynthetic ICMs in three phylogenetically disparate alphaproteobacteria, *R. sphaeroides (Rhodobacterales), Rhodospirillum rubrum (Rhodospirillales)*, and *R. palustris (Rhizobiales)* ^55,56,64,65^, as indicated by the proteomes of isolated ICMs. Third, *R. sphaeroides’* Mic60 is capable of binding and tubulating membranes *in vitro*, and its heterologous overexpression induces ICM-like structures in *E. coli* ^27^. Fourth, both *mic60* and *orf52* have paralogs in the magnetosome gene island of the alphaproteobacterial genus *Magnetospirillum* of the order *Rhodospirillales* ^19^ whose disruption leads to fewer and larger magnetosomes ^66,67^. Multiple sources of evidence thus support the notion that the function of Mic60 has been conserved in alphaproteobacteria.

What is the precise role of Mic60 in ICM development? Mic60 likely introduces curvature at ICM junctions through an amphipathic helix that is conserved between mitochondria and alphaproteobacteria (see Fig. 2). That alphaproteobacterial Mic60 binds to and bend membranes has already previously been shown *in vitro* ^27^. Moreover, alphaproteobacterial Mic60 most likely interacts with BamA, the central subunit of the BAM complex and homolog of Sam50, as shown in this study (Fig. 5). In mitochondria, Sam50 interacts with Mic60 through its intermembrane space-protruding POTRA domains, and it is likely that the same type of interaction occurs in the periplasm of alphaproteobacteria. This ancient protein-protein interaction suggests that, in addition to aiding the formation of ICM junctions, alphaproteobacterial Mic60 is also involved in the formation of contact sites that anchor ICMs to the alphaproteobacterial envelope. The formation of crista junctions and contact sites by Mic60 are aided by Mic19 in mitochondria ^40,42^, but there is no evidence for a Mic19-like protein in alphaproteobacteria; our pull-downs did not reveal proteins with Coiled-Coil-Helix-Coiled-Coil-Helix (CHCH) motifs. In summary, it appears that both the formation of contact sites and Crista/ICM junctions have been conserved in alphaproteobacteria and mitochondria.

The larger macromolecular complex that Mic60 is part of in *R. sphaeroides* is reminiscent of the extended interaction network, ERMIONE, that in *S. cerevisiae* mitochondria involves Mic60 ^68^. In *S. cerevisiae*, the ER–mitochondria organizing network, or ERMIONE, plays a major role in mitochondrial biogenesis by connecting the ER–mitochondria encounter structure (ERMES) to MICOS through the outer membrane SAM and TOM complexes ^11,68^. In *H. sapiens*, MICOS and SAM interact stably to form the Mitochondrial Intermembrane space Bridging complex (MIB) complex ^69^ It is possible that BamA, through its interactions with Mic60, Orf52, OM Efflux, and RSP_7515, serves as the hub of a larger protein-interaction network, that, for example, facilitates lipid transfer and protein export to the outer membrane. In the model alphaproteobacterium *Caulobacter crescentus*, the BAM complex has been shown to have a modular structure (including a ~300 kDa subcomplex) and interact with Pal, a lipoprotein that serves as an anchor to the peptidoglycan layer of the cell wall ^70,71^. In addition, contact sites may help to stabilize ICMs by providing an anchor to the OM at points of high membrane curvature, i.e., ICM junctions ^72^. This may contribute to the biogenesis of ICMs by ensuring their continuity with the CM where protein complex subunits may first be inserted and assembled to give rise to UPBs.

The evidence for the structural and functional conservation of alphaproteobacterial Mic60 relative to its mitochondrial homolog is most compatible with an evolutionary scenario in which ICMs and cristae are homologous. This view implies that cristae most likely evolved from the ICMs developed by the last common ancestor of mitochondria and its sister group, the *Alphaproteobacteria;* cristae thus have a pre-endosymbiotic origin. If this hypothesis turns out to be correct, bioenergetic ICMs might have pre-adapted the first mitochondrial ancestor to become an efficient bioenergetic or respiratory organelle ^26^ The widespread but sporadic phylogenetic distribution of ICMs across the *Alphaproteobacteria* is then most likely explained by multiple independent losses. This is conceivable as both cristae and ICMs are known to have been lost repeatedly as a result of physiological specialization to different environments (e.g., transitions to anaerobiosis in mitochondria or to heterotrophy in photosynthetic bacteria). However, although many alphaproteobacteria may be capable of developing ICMs under certain environmental conditions (e.g., aerobic phototrophs or *C. crescentus* under low oxygen conditions ^73^), some probably never develop ICMs despite having Mic60 homologs. In these alphaproteobacteria, Mic60 might play a more general function, e.g., contact site formation by interacting with BamA for lipid transfer or protein export, that is still required in the absence of ICMs. This more general function might have been ancestral to Mic60 and agrees with the observation that a distant homologue of Mic60 that lacks the conserved C-terminal mitofilin domain, namely HemX, is restricted to and widespread in the *Gammaproteobacteria* ^20^. It is thus conceivable that the signature mitofilin domain first evolved in a HemX-like protein and that this coincided with the origin of ICMs in a common ancestor of alphaproteobacteria and mitochondria.

Future studies are required to elucidate the molecular mechanisms by which alphaproteobacterial Mic60 interacts with BamA, and determine whether alphaproteobacterial Mic60 forms homodimers and homotetramers as recently reported for fungal Mic60 ^42^. More generally, efforts focused on phylogenetically disparate alphaproteobacteria with and without ICMs will shed light on the functions and mechanisms of Mic60 in prokaryotes. Altogether, the structural and functional conservation of alphaproteobacterial Mic60 shown here, suggests a role of this protein in curving membranes at ICM junctions and making contact sites at envelopes. Iit is therefore probable that the mitochondrial ancestor was an ICM-bearing alphaproteobacterium.

## MATERIALS AND METHODS

### Bacterial strains, media and growth conditions

*R. palustris* TIE-1 and *E. coli* BW29427-λpir-RP4 were kindly provided by Dianne K. Newman (California Institute of Technology) ^74^ *R. sphaeroides* 2.4.1, *E. coli* S17-1-λpir, and *E. coli* DH5α-λpir were kindly provided by Jeanette Johnson-Beatty (University of British Columbia). *R. palustris* TIE-1 strain was grown both chemo- and photo-heterotrophically at 30 °C in YPS rich medium (Jiao et al., 2005) unless otherwise noted. *R. palustris* TIE-1 was grown photoheterotrophically at 30 °C on FEM minimal medium ^75^. *R. sphaeroides* 2.4.1 strain was grown chemoheterotrophically at 30 °C in LB or RLB rich media ^76^, unless otherwise noted, and photoheterotrophically at 30 °C on RCVBN minimal medium ^77^ Table S1 lists all strains used this study.

For growth analysis under photoheterotrophic conditions, *R. sphaeroides* and *R. palustris* mutants and WT strains were grown in front to three incandescent 40 W lightbulbs at a temperature of 29 °C, with the distances of the culture tubes from the light sources adjusted to allow the appropriate light intensities and temperature. The light intensities were measured to be ~200 μmol photons m-^2^ s^-1^ for high light and ~10 μmol photons m-^2^ s^-1^ for low light. The latter was achieved by placing the two layers of approximately ~ 50 % neutral density filter sheeting between the light source and culture tubes. Prior to growth analysis, cells were grown in in rich media in triplicate (*R. sphaeroides*, RLB without antibiotics; *R. palustris*, YPS no antibiotic for WT, 400 μg/ml kanamycin for mutants) until mid-log phase. At mid-log phase equivalent amounts of cells (turbidity x volume = 20, i.e., if the turbidity value was equal to 5.10, then 3.92 ml of cells were decanted) were removed from the growth tubes into 15 ml Falcon tubes and pelleted by centrifugation. The pellet was washed once in minimal media and resuspended in 1 ml of the required minimal media (*R. sphaeroides* 10 ml RCVBN + 8 ml RLB and *R. palustris* FEM). A 150 μl volume was of each cell line suspension was inoculated in triplicate; this was time point 0. Turbidity measurements were performed using a McFarland Densitometer DEN-1B that measures at λ = 565 ±15 nm. Turbidity measurements were then taken every 6 h for a total experiment length of 72 h. For analysis of chemoheterotrophic growth, the same approach was made except the cell cultures were grown in the dark in well-aerated flasks that were under constant agitation to ensure gas exchange. Whole-cell absorption spectra of these cells were measured to verify the absence of LH1 and LH2 complexes under these conditions (Fig S3D). Spectra were recorded from bacterial strains grown to late-log phase at all photoheterotrophic and chemoheterotrophic conditions after resuspension in MES buffer pH 6.8 resuspended to OD ~6,5. The absorption spectra were measured at 0.5 nm intervals using a Shimadzu UV-Vis-NIR UV2600 spectrophotometer equipped with an integrating sphere. The resulting spectra were normalised to OD = 1 at the BChl Qx (~ 590 nm) peak for comparison.

### DNA methods and plasmid construction

Total genomic DNA (gDNA) was extracted from *R. sphaeroides* and *R. palustris* strains using the ZR Bacterial DNA Miniprep Kit (Zymo Research) with a BIO101/Savant FastPrep FP120 high-speed bead beater and a 30 min incubation at 60°C with 20 μl of proteinase K (20 mg/mL), or the Epicentre MasterPure DNA Purification Kit (Epicentre Biotechnologies). All plasmid constructs were built with the Gibson Assembly Master Mix or the NEB Builder HiFi DNA Assembly (both New England Biolabs). Plasmids were isolated using the AxyPrep Plasmid Miniprep Kit (Axygen). All plasmid constructs were confirmed by Sanger sequencing using both forward and reverse primers. Table S2 lists all plasmids used. Table S3 lists all primers used in this study.

### Construction of *R. sphaeroides* and *R. palustris* knockout strains

To knock out genes in *R. sphaeroides*, a knockout construct was assembled into the suicide plasmid vector pZDJ ^78^ with either the Gibson Assembly Cloning Kit or the NEBuilder HiFi DNA Assembly Master Mix (both New England Biolabs). The antibiotic resistance cassette flanked by flippase recognition target (FRT) sites used to interrupt the genes to be knocked out corresponds to that used in the Keio collection ^79^. The resulting suicide vector with the knockout construct was cloned into *E. coli* S17-1 λ-pir-RP4, which can replicate the suicide plasmid. This strain was then conjugated with *R. sphaeroides 2.4.1*. Briefly, donor and recipient cells grown to stationary phase were mixed in a 1:2 volume ratio and pelleted by centrifugation for 1 min at 4,500-6,000 × g, and then washed twice with antibiotic-free RCVBN minimal medium. After the last wash, the pellet was resuspended in 50 μl of RCVBN, and 10 μl aliquots were spotted onto antibiotic-free RCVBN solid medium. The plates were incubated at 30° C overnight to allow conjugation to take place. Afterwards, an emulsion was made from the several inoculation spots and streaked onto antibiotic-containing RCVBN+Gm solid medium. *R. sphaeroides* exconjugants were then successively streaked onto new LB+Gm solid medium until no *E. coli* S17-1 λ-pir-RP4 contamination remained. The resulting pure *R. sphaeroides* exconjugants contained the pZDJ plasmid, with the knockout gene construct, integrated in the chromosome by a first crossing over (recombination) event. To induce a second crossing over to excise the pZDJ suicide plasmid from the host chromosome, colonies were picked and grown on liquid LB with no antibiotic selection until late stationary phase (about 2-3 days). The cultures were then streaked on LB+10% sucrose solid medium, which allows for counter-selecting of those colonies that have lost the integrated pZDJ plasmid. The counter-selection relies on the *sacB* gene carried by the suicide plasmid ^50^. In order to induce a third crossing over between the FRT sites of the kanamycin cassette, cells were grown in liquid LB without any antibiotic selection until late stationary phase and then streaked on LB solid medium. Resultant colonies were then screened through PCR assays.

To make knockout strains for *R. palustris*, the same general protocol used for *R. sphaeroides* was followed ^80^. The suicide plasmid used for R. palustris was pJQ200SK ^81^ and the host strain was *E. coli* BW29427-λpir-RP4, which requires 300 μM diaminopimelic acid for growth. This auxotrophy allows to easily remove the plasmid-donor bacterium from the medium after conjugation.

### Construction of *R. sphaeroides* and *R. palustris* overexpression strains

To create Mic60- and Orf52-overexpressing strains, the inducible expression plasmids for *R. sphaeroides* pIND4 -Km and pIND4-Gm were kindly provided by Judith P. Armitage (University of Oxford), and Alexander Westbye (University of British Columbia), respectively. The inducible expression plasmid pSRK was used in *R. palustris*. The coding sequences of the *mic60* and *orf52* genes of *R. sphaeroides* and *R. palustris* were amplified by PCR and assembled into the expression vectors pIND4 and pSRK, respectively, with the NEBuilder HiFi DNA Assembly Master Mix (New England Biolabs). The *mic60* and *orf52* homologs were cloned into kanamycin resistance-conferring plasmids and gentamycin resistance-conferring plasmids, respectively. To insert the C-terminal 6xHis tag as part of the coding sequences of *mic60* and *orf52* the BamHI and BglII restriction sites in pIND4 were used, or the tag was incorporated as part of the primer used to amplify the targets in *R. palustris*. The assembled plasmids were transformed into suitable conjugative *E. coli* hosts, and conjugation assays with recipient *R. sphaeroides* and *R. palustris* strains were done as described above. After conjugation, the exconjugants were repeatedly streaked onto antibiotic-containing plates to remove the plasmid-donor bacterium. The resulting overexpression strains were further verified by RT qPCR.

### RT qPCR

*R. sphaeroides* and *R. palustris* strains were grown in triplicate until mid-log phase as described for growth analysis, after which 1 mM IPTG was added, and the cells allowed to grow for 7 h 20 min for *R. sphaeroides* and 4 h 30 min for *R. palustris;* cultures were grown in parallel in the absence of IPTG. Then 1 ml of cell suspension was centrifuged and the pellet resuspended in 1 ml of PGTX nucleic acid extraction buffer ^82^. The samples were then flash frozen and stored at −20 °C until the PGTX-mediated nucleic acid extraction procedure, essentially following the classical phenol/chloroform extraction method (Pinto et al., 2009). Briefly, isolated total nucleic acid concentration was measured using a NanoDrop DeNovix DS-11 (Thermo Fisher) and diluted to 100 ng/μL in 50 μl DNase/RNase-free H_2_O. The digestion of gDNA was performed by addition of DNase I (Qiagen) and the concentration of total RNA was measured using the Qubit 2.0 Fluorometer (Thermo Fisher). A total of 100 ng RNA was subsequently reverse transcribed using the Transcriptor First Strand cDNASynthesis Kit (Roche) at 55 °C for cDNA synthesis. Quantitative PCR (qPCR) was performed in triplicates in a CFX96 qPCR cycler (Bio-Rad) in 20 μl reactions containing 1x PowerUp Sybr Green master mix (Applied Biosystems, USA), 8 pmol of each primer, and 10 μg cDNA with the following program: initial denaturation 10 min at 95 °C; 45 cycles of 15 s at 95 °C and 30 s at 60 °C. Primers annealing to *mic60, orf52*, and the *rpoZ* cDNAs used for qPCR are listed in Table S3. Cycle threshold values were automatically computed with the CFX Maestro software (Bio-Rad). Additionally, non-reverse transcribed RNA sample was used as a control to verify complete degradation of genomic DNA. Relative abundance of transcripts in cells grown in the presence of IPTG compared to those grown without the inducing agent was calculated using the 2^-ΔΔCT^ method ^83^; the unaffected *rpoZ* housekeeping gene cDNA was used to normalize the *mic60* and *orf52* values.

### Transmission electron microscopy

Both classical TEM and electron tomography on 80 nm-thick sections were done as previously described (Cadena et al., 2021; Kaurov et al., 2018). Scoring of *R. palustris* ICM area/TEM area and *R. sphaeroides* tubules/cell per section were performed blinded on images obtained using JEOL 1010 TEM operating at an accelerating voltage of 80 kV and equipped with a MegaView III CCD camera (SIS). Measurement of *R. palustris* ICM and total cell areas was done using Image J software ^84^ by tracing along the outermost electron dense membranes of the ICM network and the outer membrane, respectively. The occurrence of tubule-like and branching ICMs in *R. sphaeroides* was counted on 20-21 images that were taken at 40,000x magnification (2.56 nm/pixel) at random parts of the grid as before (Kaurov et al., 2018). These images were mixed and randomized prior to blind scoring.

Electron tomograms were collected at a range of ±65° with tilt 1° steps using the JEOL 2100F TEM working at 200 kV, equipped with Gatan camera K2 Summit and controlled with SerialEM automated acquisition software ^85^ IMOD software ^86^ was used for tomogram reconstruction and generating 3D models by segmentation.

### SDS PAGE and Western blotting

Bacterial lysates were separated on a Bolt 4-12% Bis-Tris Plus gel (Invitrogen), blotted onto a PVDF membrane (Amersham), blocked in 5% low-fat, powdered milk (w/v) in phosphate buffered saline with 0.1% Tween 20 (v/v) (PBS-T), and probed with 6x-His Tag Monoclonal Antibody (HIS.H8) (Thermo Fisher Scientific #MA1-21315) diluted in 5% milk in PBS-T (1:500). This was followed by incubation with secondary HRP-conjugated anti-mouse antibody (1:2000; Bio-Rad). Proteins were visualized using the Pierce ECL system (Genetica/Bio-Rad) on a ChemiDoc imager (Bio-Rad).

### Isolation of ICMs via high-pressure homogenization and light spectroscopy

*R. sphaeroides* strains were cultured identically in C-succinate media ^87^ in flat, glass bottles at 28-30°C under anaerobic conditions. Illumination was provided by one incandescent 40W bulb providing ~10 μmol photons m^-2^ s^-1^. The cells were fully adapted to the incident light-intensity and harvested by centrifugation once they had reached mid-log phase. Cell pellets were washed in 20 mM MES, 100 mM KCl, pH 6.8, then flash frozen and stored at −80 °C until required

The cell pellets were re-suspended in 20 mM Tris.Cl, pH 8.0 and homogenised thoroughly with a few grains of DNAse I (Qiagen) and a few mg of MgCl_2_. The cells were broken by passage three times through an Emulsiflex-C5 cell disrupter (Avestin). The ruptured cell solution was first subjected to a low-speed centrifugation step (10 min, 10,000 × *g*, 4°C) to remove any unbroken cells. The decanted supernatant was ultra-centrifuged (120 min, 180,000 × *g*, 4°C), after which the supernatant was discarded. The resulting chromatophore pellet was gently re-suspended in 20 mM Tris-HCl pH 8.0 before the optical density was adjusted to 10 cm^-1^ at the Qx absorption maximum (~590 nm) using a Shimadzu UV-Vis-NIR UV2600 spectrophotometer equipped with an integrating sphere.

### Blue Native PAGE

BN-PAGE of isolated ICMs was adapted from published protocols (Cadena et al., 2021). Briefly, 1 mg of total protein was resuspended in 100 μl NativePAGE sample buffer (Invitrogen), lysed with 1.5% digitonin (v/v) and 0.1% Triton X-100 (v/v) for 1 h on ice then cleared by centrifugation (22,000 × g, 20 min, 4 °C). Subsequently, 5% Coomassie brilliant blue G-250 was added before loading ~100 μg on a 3-12% Bis-Tris BNE gel (Invitrogen). After electrophoresis (2.5 hours, 150 V, 4 °C), the gel was blotted onto a PVDF membrane (Amersham) and probed as described above.

### Affinity purification

Affinity purification (AP) of tagged proteins from 1 mg isolated ICMs were solubilized in IPP50 buffer (50 mM KCl, 20 mM Tris-HCl pH 7.7, 3 mM MgCl_2_, 10% glycerol, 1 mM phenylmethanesulfonyl fluoride, complete EDTA free protease inhibitor cocktail (Roche) supplemented with 1.5% digitonin (v/v) and 0.1% Triton X-100 (v/v) for 1 h on ice. After centrifugation (22,000 ×g, 20 min, 4 °C) the supernatant was added to 2.0 mg of cobalt conjugated Dynabeads (Thermo Fisher) to capture the His-tag. The Dynabeads were pre-washed in 300 μl of IPP50 + 1.5% digitonin for 5 min at RT. The solubilized ICMs were rotated with beads for 90 min at 4 °C. After removal of the flow-through, the beads were washed three times in IPP50 + 1.5% digitonin. Prior to eluting, the beads were transferred into a new tube. Elution was done with 300 mM imidazole in IPP50 for 15 min at RT and shaking at 1000 rpm. The elutes were further processed for LC-MS^2^ analysis or resolved by SDS-PAGE. APs were performed in triplicate.

### Protein preparation and mass spectroscopy

Individual bands containing proteins of interest were excised from Coomasie stained SDS PAGE gel using a razor blade and cut into small pieces (~1 mm^3^). Bands were destained by sonication for 30 min in 50% acetonitrile (ACN) and 50 mM ammonium bicarbonate (ABC). After destaining, gels were dried in ACN. Disulfide bonds were reduced using 10mm DTT in 100mM ABC at 60°C for 30 min. After that, samples were again dried with ACN and free cysteine residues were blocked using 55 mM iodoacetamide in 100 mM ABC for 10 min at room temperature in the dark. Samples were dried thoroughly and then digestion buffer (10% ACN, 40 mM ABC and 13 ng/μl trypsin) was added to cover gel pieces. Proteins were digested at 37 °C overnight. After digestion, 150 μl of 50% ACN with 0,5% formic acid was added and sonicated for 30 min. Supernatant containing peptides was transferred to a new microcentrifuge tube and another 150 μl of elution solution was added and sonicated for 30 min. This solution was removed, combined with the previous solution and dried by SpeedVac. Dried peptides were reconstitued in 2% ACN with 0,1% TFA and injected into Ultimate 3000 Nano LC coupled to Orbitrap Fusion.

Eluates of co-AP proteins and thin-sliced BN-PAGE gels were processed for MS analysis as described elsewhere (Cadena et al., 2021). In brief, eluate samples were resuspended in 100 mM tetraethylammonium bromide containing 2% sodium deoxycholate. Cysteines were reduced with 10 mM tris(2-carboxyethyl)phosphine and subsequently cleaved with 1 μg trypsin overnight at 37 °C. After digestion, 1% trifluoroacetic acid (TFA) was added to wash twice and eluates were resuspended in 20 μl TFA per μg of protein. A nano reversed-phased column (EASY-Spray column, 50 cm x 75 μm inner diameter, PepMap C18, 2 μm particles, 100 Å pore size) was used for LC/MS analysis. Mobile phase buffer A consisted of water and 0.1% formic acid. Mobile phase D consisted of acetonitrile and 0.1% formic acid. Samples were loaded onto the trap column (Acclaim PepMap300, C18, 5 μm, 300 Å pore size, 300 μm x 5 mm) at a flow rate of 15 μl/min. The loading buffer consisted of water, 2% acetonitrile, and 0.1% TFA. Peptides were eluted using a Mobile phase B gradient from 2% to 40% over 60 min at a flow rate of 300 ml/min. The peptide cations eluted were converted to gas-phase ions via electrospray ionization and analyzed on a Thermo Orbitrap Fusion (Q-OT-qIT, Thermo Fisher). Full MS spectra were acquired in the Orbitrap with a mass range of 350-1, 400 *m/z*, at a resolution of 120,000 at 200 *m/z* and with a maximum injection time of 50 ms. Tandem MS was performed by isolation at 1,5 Th with the quadrupole, high-energy collisional dissociation (HCS) fragmentation with normalized collision energy of 30, and rapid scan MS analysis in the ion trap. The MS/MS ion count target was set to 10^4^ and the max infection time at 35 ms. Only those precursors with a charge state of 2-6 were sampled. The dynamic exclusion duration was set to 45 s with a 10 ppm tolerance around the selected precursor and its isotopes. Monoisotopic precursor selection was on with a top speed mode of 2 s cycles.

### Analysis of mass spectroscopy peptides

Label-free quantification of the data were analyzed using the MaxQuant software (version 1.6.2.1) ^88^. The false discovery rates for peptides and for proteins was set to 1% with a specified minimum peptide length of seven amino acids. The Andromeda search engine was used for the MS/MS spectra against the *R. sphaeroides* 2.4.1 predicted proteome (ASM1290v2) downloaded from NCBI GenBank on October 2020. Enzyme specificity was set to C-terminal Arg and Lys, alongside for cleavage at proline bonds with a maximum of two missed cleavages. Dithiomethylation of cysteine was selected as a fixed modification with N-terminal protein acetylation and methionine oxidation as variable modifications. The ‘match between runs’ feature in MaxQuant was used to transfer identification to other LC-MS/MS runs based on mass and retention time with a maximum deviation of 0.7 min. Quantifications were performed using a label-free algorithm as previously described (Cox et al., 2014). Data analysis was performed using Perseus software (version 1.6.1.3). Eluate co-AP proteins identified with a mean Log2 ratio (protein_His_/WT) >3-fold change and having a Andromeda confidence score >100 (Cox et al., 2011) in three independent biological replicates were analyzed, while gel-slice proteins identified with a mean Log2 transformed LFQ score >23 and present in four biological replicates were analyzed.

### Quantification and statistical analysis

Statistical significance as determined by unpaired t-test using the GraphPad Prism 7 are reported in the figures and legends.

### Data Availability

The LC-MS/MS data have been deposited to the ProteomeXchange Consortium (http://www.proteomexchange.org) via the PRIDE partner repository with the data set identifier PXD032747.

## ACKNOWLEDGEMENTS

We would like to dedicate this work to Thomas Cavalier-Smith whose large-scale synthetic work on evolutionary cell biology stimulated thinking on this topic by SAM-G many years ago. We thank Karel Harant and Pavel Talacko (Charles University, Prague) for performing LC-MS analysis and Michala Boudová (Center Algatech) for technical assistance. SAM-G is supported by an EMBO Postdoctoral Fellowship (ALTF 21-2020). MML was supported by a Nova Scotia Health Research Foundation (NSHRF) Scotia Scholarship 2012-8781. This work was also supported by the Czech Science Foundation grants 20-23513S to HH, 22-01-26S to JL and 19-28778X to MK, ERD Fund (003/0000441) to TB, as well as the Czech Ministry of Education grant OPVVV16_019/0000759 and Czech BioImaging grant LM2015062. AJR and JTB were supported by Natural Sciences and Engineering Research Council of Canada (grants RGPIN-2022-05430 and RGPIN-2018-08398, respectively).

## Supplemental Material

**Figure S1. AlphaFold2 predictions of mitochondrial and alphaproteobacterial Mic60 homologs. A.** Predicted tertiary structure of the Mic60 homolog of the yeast *L. thermotolerans*. α-helices are colored according to domain (mitofilin in red, LBS1 and LBS2 in blue, middle coiled coils in green, and transmembrane segment in orange) and follow the coordinates predicted by JPred4. **B**. Predicted tertiary structure of the Mic60 homolog of the yeast *L. thermotolerans* colored by the pLDDT scores, which denote confidence of predicted structure. The long α1C helix (207-382) whose structure was experimentally resolved by Bock-Bierbaum *et al*. (2022) is indicated. **C.** Predicted tertiary structures of the Mic60 homolog of the alphaproteobacterium *R. sphaeroides*. **D**. Predicted tertiary structure of the Mic60 homolog of the alphaproteobacterium *R. sphaeroides* colored by the pLDDT scores.

**Figure S2. Verification of *mic60* and *orf52* knockout strains and IPTG-induced overexpression of Mic60 and Orf52. A-B.** PCR assays confirm the disruption of the *mic60* and *orf52* genes in *R. palustris* (**A**) and *R. sphaeroides* (**B**). Genomic contexts of the relevant loci for the WT, Δ*mic60* and Δ*orf52* strains shown on top, with primer pairs and their expected amplicon sizes shown below each schematic gene arrangement. Lower panel shows each PCR amplicon from each strain (labelled above gel) after agarose gel electrophoresis. Size markers shown either to the left or right of the gel. (**C**) Real time PCR showing relative abundancies of Mic60 and Orf52 mRNAs in *R. palustris* (left) and *R. sphaeroides* (right) strains grown in the presence of the expression induction agent IPTG relative to the same strains grown without IPTG. Error bars show standard deviation from three replicates of assayed induced and non-induced cells.

**Figure S3. Chemoheterotrophic growth and absorption spectra of *mic60* and *orf52* knockout strains. A**. Growth curves of *R. palustris* (left) and *R. sphaeroides* (right) under chemoheterotrophic conditions in the presence of malate to feed the respiratory chain. Figure labelled as in Figs. 3A–D. Absorption spectra of whole *R. palustris* (left) and *R. sphaeroides* (right) WT and knockout strains grown photoheterotrophically at either high light (**B**) or low light (**C**), as well as chemoheterotrophically in the dark and presence of oxygen (**D**).

**Figure S4. Quantification branched ICM occurrence in *R. sphaeroides mic60* knockout (Δ*mic60)* and Mic60 overexpression (Mic60↑) strains.** (A) Representative transmission electron micrographs of elongated (hollow arrowhead) and branched (solid arrowhead) ICMs. Imaged strain indicated in upper corner of micrographs. Scale bar, 100 nm. Scatter plots showing blind quantification of branched (B) and elongated (C) ICMs. Middle bar shows median value and whiskers denote interquartile range. Statistical significance: **, P<0.01; n.s., not significant.

**Figure S5. Proteomic analysis of isolated ICMs plus Mic60 and Orf52 interactomes. A.** Blue native gel resolved detergent-solubilized ICMs from *R. sphaeroides* WT in which the ~250 kDa band (boxed) was excised in quadruplet for subsequent MS analysis. To right of gel is a histogram of the Coomassie-stained band intensities *(I) along* the vertical axis of the run. The scissors denote the ~250 kDa band intensity signal. **B-C.** A list of all proteins, including excluded contaminants, found within the enriched protein area of the volcano plots in in Figure 4A–B for Mic60 (B) and Orf52 (C). Columns as described in legend of Figure 5C–D. Note the presence of likely contaminant found in both APs, PAS-fold containing histidine kinase (PAS), which is a large protein amenable to LC-MS/MS in contrast to RSP_7517 and not found in any of our requisite proteomes. **D.** Kyle and Doolittle hydropathy plot of RSP_7517, whose amino acid sequence is given below. Predicted transmembrane domain shaded and a potential oligomerization motif AxxxA underlined. Lysine (K) residues recognized by trypsin protease are in bold.

**Dataset 1.** List of proteins found within Blue Native gel slices as indicated in Figures 3C and S4A in quadruplicates.

**Dataset 2.** List of all proteins found in the Orf52 and Mic60 interactomes in comparison to WT negative controls by MS of AP eluates. Related to Figure 4A–B.

**Table S1.** List of strains used in this study.

**Table S2.** List of plasmids used in this study.

**Table S3.** List of primers and their sequences used in this study.

**Movie S1.** 3D reconstruction model of *R. sphaeroides* WT as rendered from electron tomograms shown at the start of the video. Red, ICM membranes; yellow, cytoplasmic membrane; green, outer membrane. Lower right corner, 200 nm scale bar.

**Movie S2.** 3D reconstruction model of *R. sphaeroides Δmic60* as rendered from electron tomograms shown at the start of the video. Red, ICM membranes; yellow, cytoplasmic membrane; green, outer membrane. Lower right corner, 200 nm scale bar.

**Movie S3.** 3D reconstruction model of *R. sphaeroides* Mic60↑ as rendered from electron tomograms shown at the start of the video. Red, ICM membranes; yellow, cytoplasmic membrane; green, outer membrane. Lower right corner, 200 nm scale bar.

**Movie S4.** 3D reconstruction model of *R. sphaeroides Δorf52* as rendered from electron tomograms shown at the start of the video. Red, ICM membranes; yellow, cytoplasmic membrane; green, outer membrane. Lower right corner, 200 nm scale bar.

**Movie S5.** 3D reconstruction model of *R. sphaeroides* Orf52↑ as rendered from electron tomograms shown at the start of the video. Red, ICM membranes; yellow, cytoplasmic membrane; green, outer membrane. Lower right corner, 200 nm scale bar.

