## Supplementary Figures for "The development of intracytoplasmic membranes in alphaproteobacteria involves the conserved mitochondrial crista-developing protein Mic60"

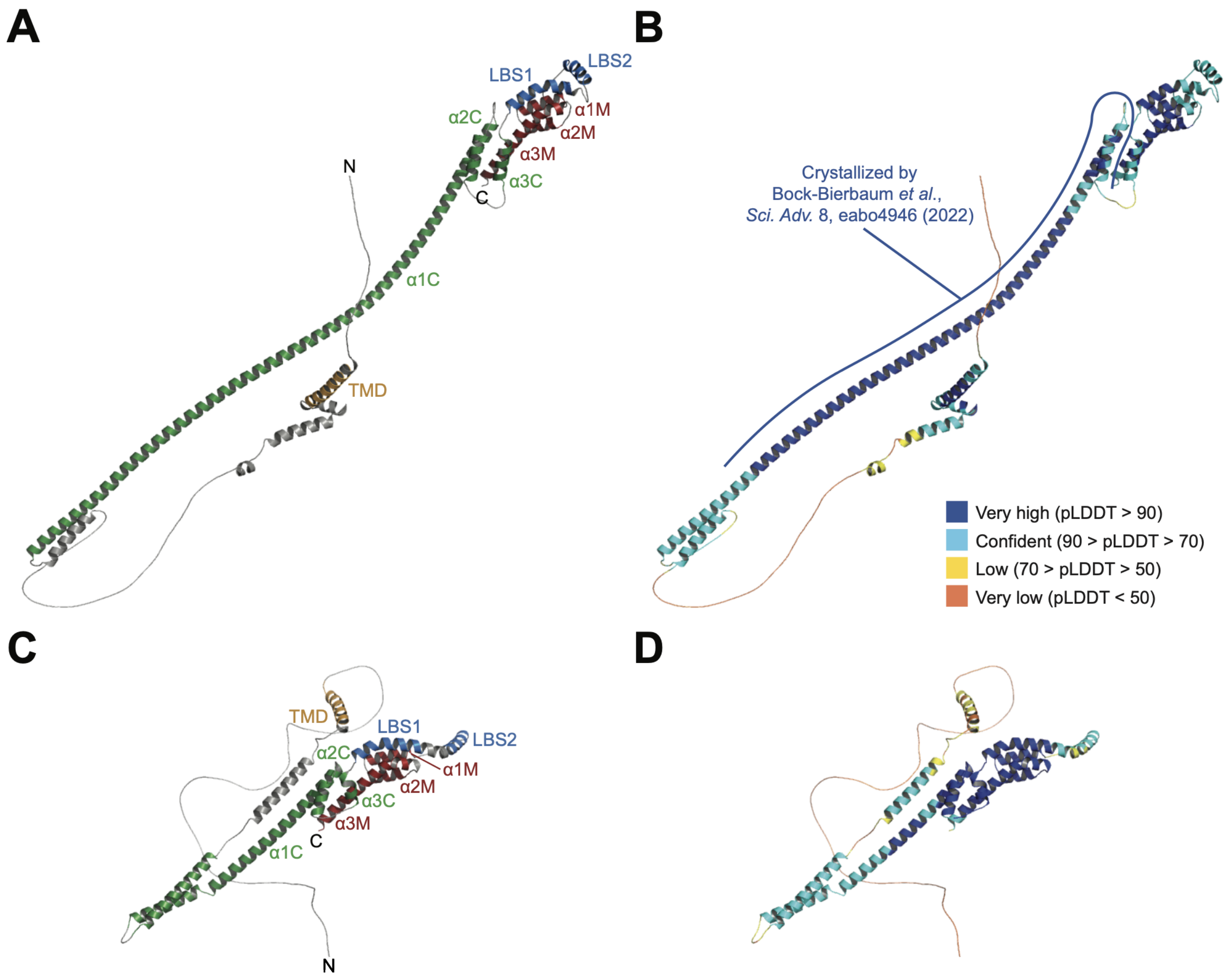


**Figure S1. AlphaFold2 predictions of mitochondrial and alphaproteobacterial Mic60 homologs.** (A) Predicted tertiary structure of the Mic60 homolog of the yeast *L. thermotolerans*. α-helices are colored according to domain (mitofilin in red, LBS1 and LBS2 in blue, middle coiled coils in green, and transmembrane segment in orange) and follow the coordinates predicted by JPred4. (B) Predicted tertiary structure of the Mic60 homolog of the yeast *L. thermotolerans* colored by the pLDDT scores, which denote confidence of predicted structure. The long α1C helix (207-382) whose structure was experimentally resolved by Bock-Bierbaum *et al.* (2022) is indicated. (C) Predicted tertiary structure of the Mic60 homolog of the alphaproteobacterium *R. sphaeroides*. (D) Predicted tertiary structure of the Mic60 homolog of the alphaproteobacterium *R. sphaeroides* colored by the pLDDT scores.

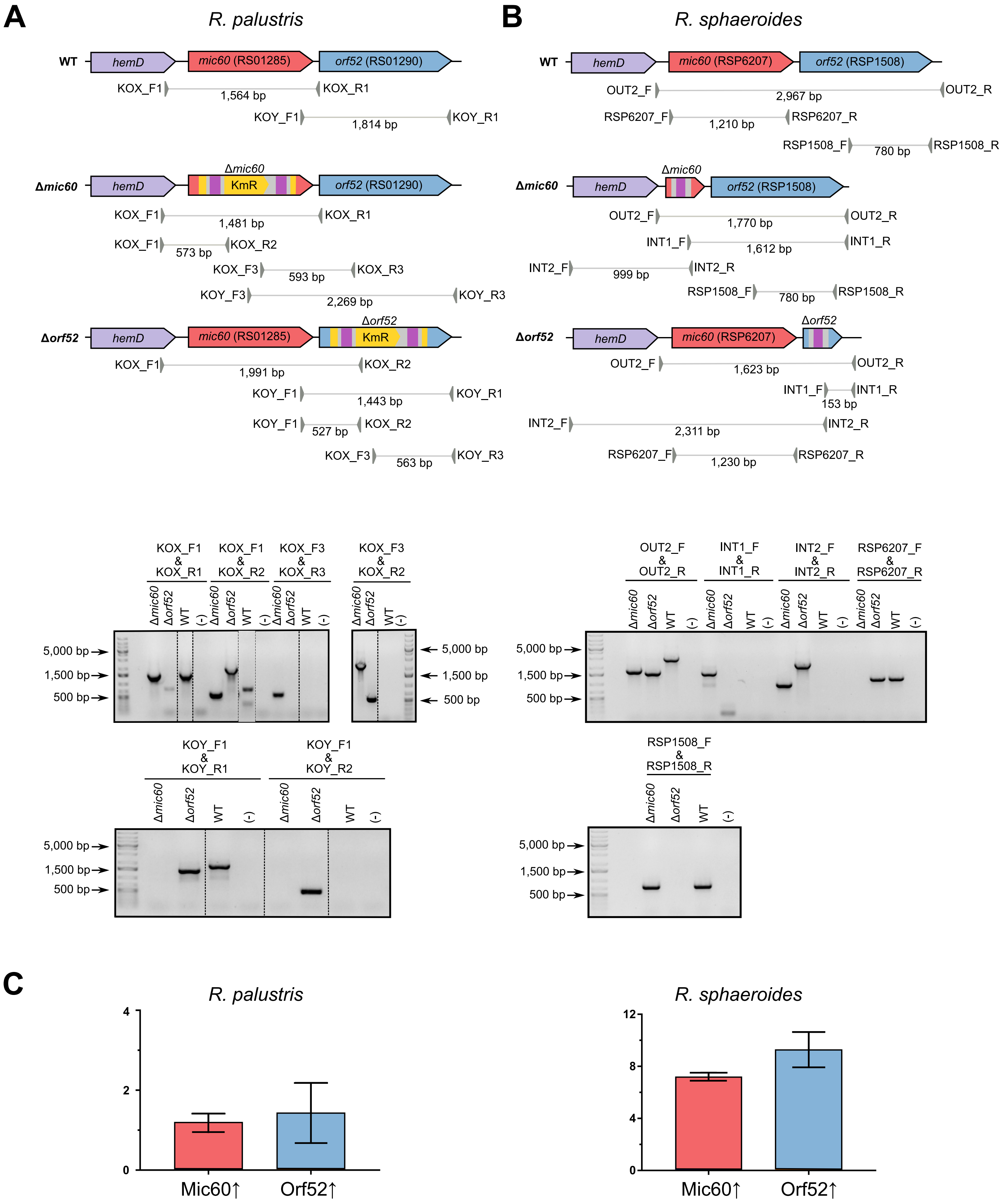
A
R. palustris
WT
hemD
mic60 (RS01285)
orf52 (RS01290)
KOX_F1
1,564 bp
KOX_R1
KOY_F1
1,814 bp
KOY_R1
Δmic60
Δmic60
KmR
KOX_F1
1,481 bp
KOX_R1
KOX_F1
573 bp
KOX_R2
KOX_F3
593 bp
KOX_R3
KOY_F3
2,269 bp
KOY_R3
Δorf52
Δorf52
KmR
KOX_F1
1,991 bp
KOX_R2
KOY_F1
1,443 bp
KOY_R1
KOY_F1
527 bp
KOX_R2
KOX_F3
563 bp
KOY_R3
B
R. sphaeroides
WT
hemD
mic60 (RSP6207)
orf52 (RSP1508)
OUT2_F
2,967 bp
OUT2_R
RSP6207_F
1,210 bp
RSP6207_R
RSP1508_F
780 bp
RSP1508_R
Δmic60
Δmic60
OUT2_F
1,770 bp
OUT2_R
INT1_F
1,612 bp
INT1_R
INT2_F
999 bp
INT2_R
RSP1508_F
780 bp
RSP1508_R
Δorf52
Δorf52
OUT2_F
1,623 bp
OUT2_R
INT1_F
153 bp
INT1_R
INT2_F
2,311 bp
INT2_R
RSP6207_F
1,230 bp
RSP6207_R
KOX_F1 & KOX_R1
KOX_F1 & KOX_R2
KOX_F3 & KOX_R3
KOX_F3 & KOX_R2
Δmic60 Δorf52 WT (-)
5,000 bp
1,500 bp
500 bp
KOY_F1 & KOY_R1
KOY_F1 & KOY_R2
OUT2_F & OUT2_R
INT1_F & INT1_R
INT2_F & INT2_R
RSP6207_F & RSP6207_R
RSP1508_F & RSP1508_R
C
R. palustris
0
2
4
Mic60↑
Orf52↑
R. sphaeroides
0
4
8
12
Mic60↑
Orf52↑

**Figure S2. Verification of *mic60* and *orf52* knockout strains and IPTG-induced overexpression of Mic60 and Orf52.** A-B. PCR assays confirm the disruption of the *mic60* and *orf52* genes in *R. palustris* (A) and *R. sphaeroides* (B). Genomic contexts of the relevant loci for the WT, *Δmic60* and *Δorf52* strains shown on top, with primer pairs and their expected amplicon sizes shown below each schematic gene arrangement. Lower panel shows each PCR amplicon from each strain (labelled above gel) after agarose gel electrophoresis. Size markers shown either to the left or right of the gel. (C) Real time PCR showing relative abundancies of Mic60 and Orf52 mRNAs in *R. palustris* (left) and *R. sphaeroides* (right) strains grown in the presence of the expression induction agent IPTG relative to the same strains grown without IPTG. Error bars show standard deviation from 3 replicates of assayed induced and non-induced cells.

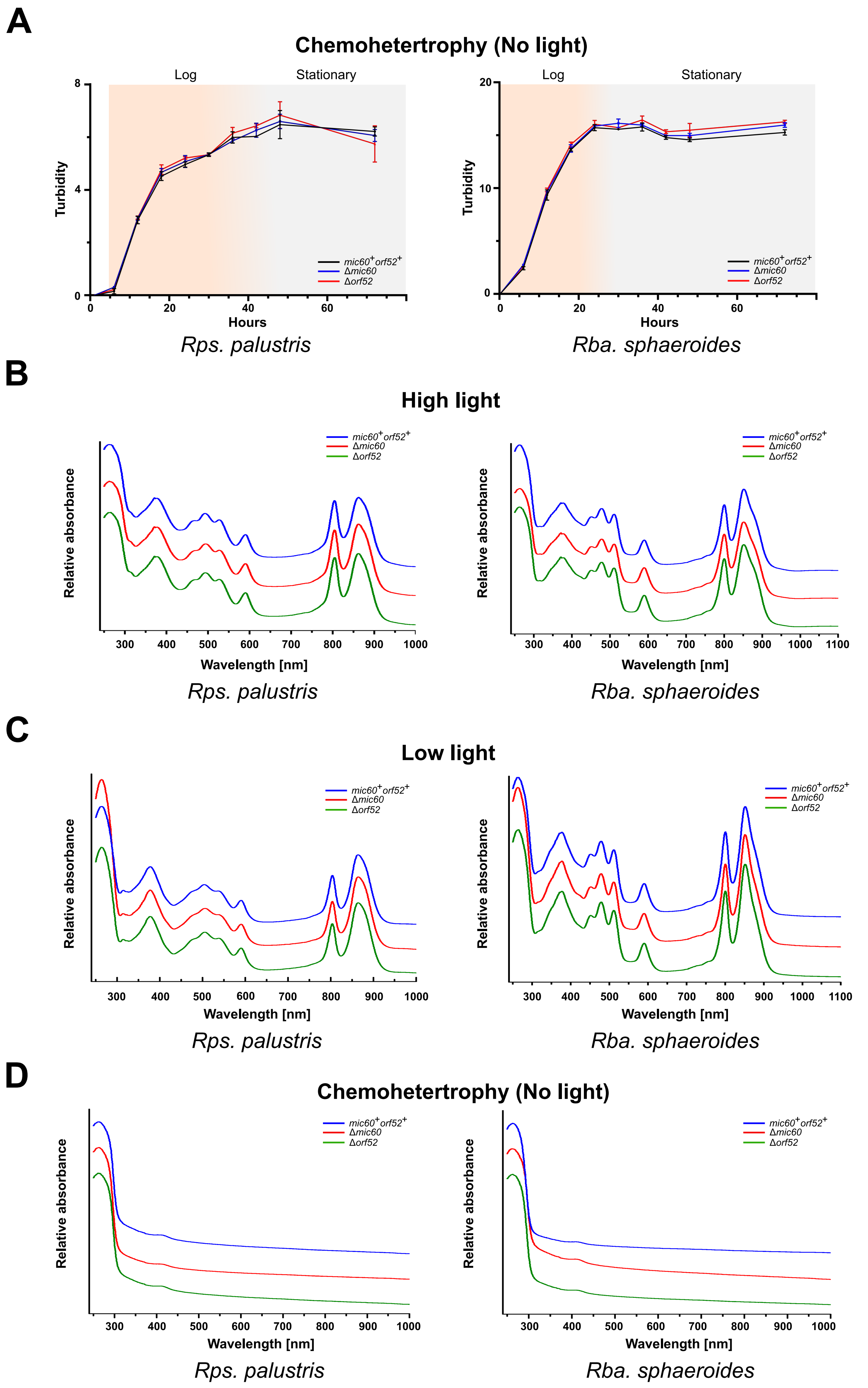


**Figure S3. Chemoheterotrophic growth and absorption spectra of *mic60* and *orf52* knockout strains.** (A) Growth curves of *R. palustris* (left) and *R. sphaeroides* (right) under chemohetertrophic conditions in the presence of malate to feed the respiratory chain. Figure labelled as in Figs. 3A-D. Absorption spectra of whole *R. palustris* (left) and *R. sphaeroides* (right) WT and knockout strains grown photoheterotrophically at either high light (B) or low light (C), as well as chemoheterotrophically in the dark and presence of oxygen (D).

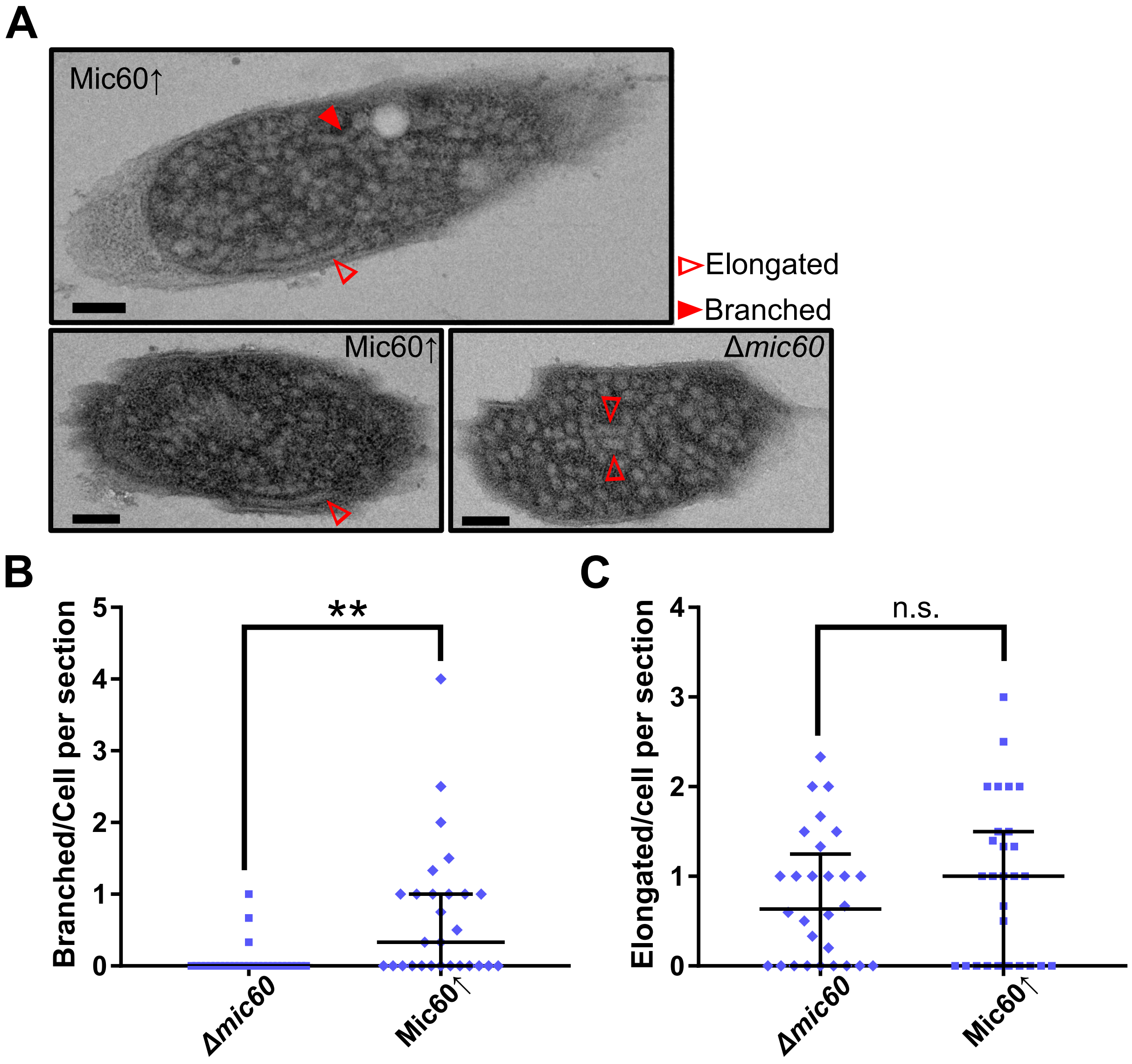


**Figure S4. Quantification branched ICM occurrence in *R. sphaeroides mic60* knockout (*Δmic60*) and Mic60 overexpression (Mic60↑) strains.** (A) Representative transmission electron micrographs of elongated (hollow arrowhead) and branched (solid arrowhead) ICMs. Imaged strain indicated in upper corner of micrographs. Scale bar, 100 nm. Scatter plots showing blind quantification of branched (B) and elongated (C) ICMs. Middle bar shows median value and whiskers denote interquartile range. Statistical significance: **, P<0.01; n.s., not significant.

**A**

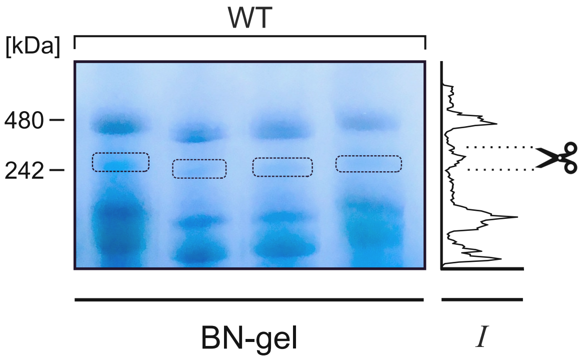


**B**

### Mic60

| Protein | Fold change | UPB | ICM | Membrane integral | 250 kDa complex | Mol. weight [kDa] |
|---|---|---|---|---|---|---|
| Mic60 | 245.13 | ◐ | ◐ * | ● | ● | 43.8 |
| PAS sensor Signal Tranduction Histidine Kinase | 9.57 | ○ | ○ | ○ | ○ | 105.0 |
| BamA | 9.09 | ● | ● | ● | ● | 88.2 |

**C**

### Orf52

| Protein | Fold change | UPB | ICM | Membrane integral | 250 kDa complex | Mol. weight [kDa] |
|---|---|---|---|---|---|---|
| Transcriptional regulator, LuxR family | 46.21 | ○ | ○ | ○ | ○ | 26.9 |
| RSP_7517 | 35.62 | ○ | ○ | ● | ◐ | 4.8 |
| Ribonucleoside-diphophate reductase class II | 31.75 | ○ | ○ | ○ | ○ | 132.0 |
| 3-oxoacyl-(acyl-carrier-protein) reductase | 26.75 | ○ | ○ | ○ | ○ | 25.0 |
| RSP_3919 (plasmid) | 23.63 | ○ | ○ | ○ | ○ | 37.5 |
| BamA | 22.38 | ● | ● | ● | ● | 88.2 |
| Transcriptional regulator, MarR family | 22.15 | ○ | ○ | ○ | ○ | 19.0 |
| OM Efflux protein | 18.52 | ● | ● | ● | ● | 49.7 |
| PAS sensor Signal Tranduction Histidine Kinase | 12.49 | ○ | ○ | ○ | ○ | 104.9 |
| Transcriptional regulator, winged helix family | 9.50 | ○ | ○ | ○ | ○ | 26.1 |
| Orf52 | 8.50 | ● | ● | ● | ● | 52.5 |
| Translation enlongation factor 2 | 8.42 | ● | ○ | ○ | ◐ | 77.7 |
| Peptidyl-tRNA hydrolase | 8.32 | ○ | ○ | ○ | ○ | 24.2 |

**D**

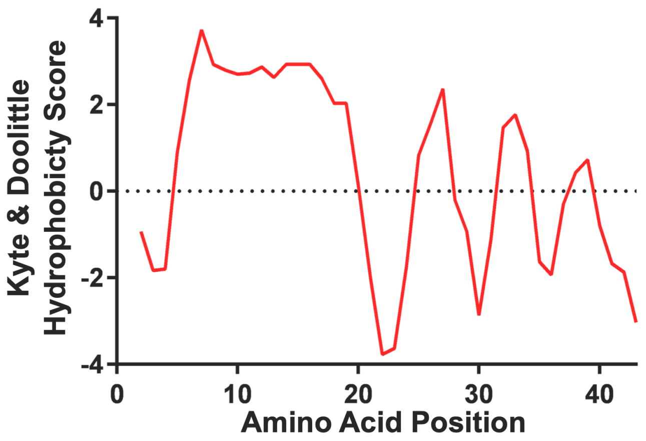


MSKSTVVFALCMVAFVAACA**KK**EEAVYVEQPAVTTEPVYTG**KYK**

**Figure S5. Proteomic analysis of isolated ICMs plus Mic60 and Orf52 interactomes.** (A) Blue native gel resolved detergent-solubilized ICMs from R. sphaeroides WT in which the ~250 kDa band (boxed) was excised in quadruplet for subsequent MS analysis. To right of gel is a histogram of the Coomassie-stained band intensities (*I*) along the vertical axis of the run. The scissors denote the ~250 kDa band intensity signal. (B-C) A list of all proteins, including excluded contaminants, found within the enriched protein area of the volcano plots in the Figure 4A-B for Mic60 (B) and Orf52 (C). Columns as described in legend of Figure 5C-D. Note the presence of likely contaminant found in both APs, PAS-fold containing histidine kinase (PAS), which is a large protein amenable to LC-MS/MS in contrast to RSP_7517 and not found in any of our requisite proteomes. (D) Kyle and Doolittle hydropathy plot of RSP_7517, whose amino acid sequence is given below. Predicted transmembrane domain shaded and a potential oligomerization motif AxxxA underlined. Lysine (K) residues recognized by trypsin protease are in bold.
