## Supplementary material for "The development of intracytoplasmic membranes in alphaproteobacteria involves the conserved mitochondrial crista-developing protein Mic60": Table S1

**Table S1.** List of strains used in this study.

| **Strain** | **Genotype** | **Source** |
| --- | --- | --- |
| *Escherichia coli* TOP10 | F^–^ *mcr*A Δ(*mrr*-*hsd*RMS-*mcr*BC) φ80*lac*ZΔM15 Δ*lac*X74 *rec*A1 *ara*D139 Δ(*ara-leu*)7697 *gal*U *gal*K λ^–^ *rps*L(Str^R^) *end*A1 *nup*G | Laboratory stock |
| *Escherichia coli* S17-1- λpir-RP4 | TpR SmR *rec*A, *thi*, *pro*, *hsd*R-M+RP4: 2-Tc:Mu: Km Tn7 λpir. | J. Thomas Beatty (University of British Columbia). |
| *Escherichia coli* DH5α | F^-^ Φ80*lac*ZΔM15 Δ(*lac*ZYA-*arg*F) U169 *rec*A1 *end*A1 *hsd*R17(r_k_^-^, m_k_^+^) *pho*A *sup*E44 *thi*-1 *gyr*A96 *rel*A1 λ^-^ | Laboratory stock |
| *Escherichia coli* DH5α-λpir | sup E44, Δ*lac*U169 (Φ*lac*ZΔM15), *rec*A1, *end*A1, *hsd*R17, thi-1, *gyr*A96, *rel*A1, λpir phage lysogen. | J. Thomas Beatty (University of British Columbia). |
| *Escherichia coli* BW29427-λpir-RP4 | RP4-2(TetS, kan1360::FRT), thrB1004, lacZ58(de).(M15), dapA1341::[erm pir+], rpsL(strR),thi-, hsdS-, pro- | K. A. Datsenko and B. L. Wanner, unpublished data. |
| *Rhodobacter sphaeroides* 2.4.1 | Wild type | J. Thomas Beatty (UBC). Originally isolated by W. R. Sistrom (University of Oregon) |
| *Rhodobacter sphaeroides* Δ*mic60* | ΔRSP6207 | This study |
| *Rhodobacter sphaeroides* Δ*orf52* | ΔRSP1508 | This study |
| *Rhodopseudomonas palustris* TIE-1 | Wild type. Isolate from Woods Hole, MA, USA. | Jiao et al., (2005) |
| *Rhodopseudomonas palustris* TIE-1 Δ*mic60* | ΔRS01285 | This study |
| *Rhodopseudomonas palustris* TIE-1 Δ*orf52* | ΔRS01290 | This study |
