## Supplementary material for "The development of intracytoplasmic membranes in alphaproteobacteria involves the conserved mitochondrial crista-developing protein Mic60": Table S2

**Table S2.** List of plasmids used in this study.

| **Plasmid** | **Description** | **Source** |
| --- | --- | --- |
| pZDJ | Suicide plasmid used to knock-out genes in *R. sphaeroides* | Brimacombe et al., (2013) |
| pZDJ-ΔRSP6207 | Suicide plasmid used to the *mic60* gene in *R. sphaeroides* | This study |
| pZDJ-ΔRSP1508 | Suicide plasmid used to knock-out the *orf52* gene in *R. sphaeroides* | This study |
| PJQ200SK | Mobilizable suicide vector; *sacB* Gm^r^ | Quandt & Hynes, (1993) |
| PJQ200SK-ΔRS01285 | Suicide plasmid used to the *mic60* gene in *R. palustris* | This study |
| PJQ200SK-ΔRS01290 | Suicide plasmid used to knock-out the *orf52* gene in *R. palustris* | This study |
| pIND4-Km | Expression plasmid used for overexpressing genes in *R. sphaeroides* | Ind et al., (2009) |
| pIND4-Km-RSP_6207-6xHis | Expression plasmid used to overexpress *mic60* in *R. sphaeroides* | This study |
| pIND4-Gm | Expression plasmid used for overexpressing genes in *R. sphaeroides* | Alexander Westbye (University of British Columbia) |
| pIND4-Gm-RSP_1508-6xHis | Expression plasmid used to overexpress *orf52* in *R. sphaeroides* | This study |
| pSRK-Km | Expression plasmid used for overexpressing genes in *R. palustris* | Khan et al., (2008) |
| pSRK-Km-RS01285-6xHis | Expression plasmid used to overexpress *mic60* in *R. palustris* | This study |
| pSRK-Gm | Expression plasmid used for overexpressing genes in *R. palustris* | Khan et al., (2008) |
| pSRK-Gm-RS01290-6xHis | Expression plasmid used to overexpress *orf52* in *R. palustris* | This study |
