## Supplementary material for "The development of intracytoplasmic membranes in alphaproteobacteria involves the conserved mitochondrial crista-developing protein Mic60": Table S3

**Table S3.** List of primers and their sequences used in this study.

| **Plasmid** | **Primer** | **Sequence (5’-3’)** | **PCR template** |
| --- | --- | --- | --- |
| pZDJ- ΔRSP6207 | 5’-MIT_T_F | TGATTACGCCAAGCTTGCATGCCTGCAGGTCGACTCTAGAAAACCGGGGTCGAGGGCTA | *R. sphaeroides* 2.4.1 WT gDNA |
|  | 5’-MIT_R | CGCCGTGCCTTCCAGTTTTT | *R. sphaeroides* 2.4.1 WT gDNA |
|  | kanFRT_T_F | CGGTCACAAAACACCTGAAAAACTGGAAGGCACGGCGATGATTCCGGGGATCCGTCGACC | pKD13 plasmid DNA or KanFRT cassette PCR product |
|  | kanFRT_T_R | CCAAAGCATGGAACCGGCCTCACTTCCCTTCCGCGGCTGCTGTAGGCTGGAGCTGCTTC | pKD13 plasmid DNA or KanFRT cassette PCR product |
|  | 3’-MIT_F | GGCCGGTTCCATGCTTTG | *R. sphaeroides* 2.4.1 WT gDNA |
|  | 3’-MIT_T_R | AACGACGGCCAGTGAATTCGAGCTCGGTACCCGGGGATCCGAGTGCTTGGGGTTGATCTC | *R. sphaeroides* 2.4.1 WT gDNA |
| pZDJ-ΔRSP1508 | 5’hemY_F | TGATTACGCCAAGCTTGCATGCCTGCAGGTCGACTCTAGACGCGCTGATGGCCGAGAT | *R. sphaeroides* 2.4.1 WT gDNA |
|  | 5’hemY_R | AATAGGAACTTCGAACTGCAGGTCGACGGATCCCCGGAATCATGGAACCGGCCTCACT | *R. sphaeroides* 2.4.1 WT gDNA |
|  | kanFRT_F | ATTCCGGGGATCCGTCGACC | pKD13 plasmid DNA or KanFRT cassette PCR product |
|  | kanFRT_R | TGTAGGCTGGAGCTGCTTC | pKD13 plasmid DNA or KanFRT cassette PCR product |
|  | 3’hemY_F | TCTAGAAAGTATAGGAACTTCGAAGCAGCTCCAGCCTACAGTTGAGCCCAGCACAAAATAGG | *R. sphaeroides* 2.4.1 WT gDNA |
|  | 3’hemY_R | AACGACGGCCAGTGAATTCGAGCTCGGTACCCGGGGATCCCCGAAATGATCCCCGACAGA | *R. sphaeroides* 2.4.1 WT gDNA |
| PJQ200SK-ΔRS01285 | KO-RpMic60-1 | AGGGAACAAAAGCTGGAGCTCCACCGCGGTGGCGGCCGCTCTAGAGCTCAGGATCAAGATCGG | *R. palustris* TIE-1 WT gDNA |
|  | KO-RpMic60-2 | GGTCGACGGATCCCCGGAATCATCCTTGGATTTTCCTCG | *R. palustris* TIE-1 WT gDNA |
|  | KO-RpMic60-3 | ACGAGGAAAATCCAAGGATGATTCCGGGGATCCGTCGA | pKD13 plasmid DNA or KanFRT cassette PCR product |
|  | KO-RpMic60-4 | TATGGCGACGGTTTTTGCAGTGTAGGCTGGAGCTGCTTC | pKD13 plasmid DNA or KanFRT cassette PCR product |
|  | KO-RpMic60-5 | CGAAGCAGCTCCAGCCTACACTGCAAAAACCGTCGCCA | *R. palustris* TIE-1 WT gDNA |
|  | KO-RpMic60-6 | AGGTCGACGGTATCGATAAGCTTGATATCGAATTCCTGCAGGACTTGGTGTCCTTGCGG | *R. palustris* TIE-1 WT gDNA |
| PJQ200SK-ΔRS01290 | KO-RpHemY-1 | AGGGAACAAAAGCTGGAGCTCCACCGCGGTGGCGGCCGCTCTAGATTCCAGGCCTTGCTGAAG3’ | *R. palustris* TIE-1 WT gDNA |
|  | KO-RpHemY-2 | GGTCGACGGATCCCCGGAATCATGGGCAGACCTATGGC | *R. palustris* TIE-1 WT gDNA |
|  | KO-RpHemY-3 | TCGCCATAGGTCTGCCCATGATTCCGGGGATCCGTCGA | pKD13 plasmid DNA or KanFRT cassette PCR product |
|  | KO-RpHemY-4 | CAGCGCGGCGGCCGGTAACCTGTAGGCTGGAGCTGCTTC | pKD13 plasmid DNA or KanFRT cassette PCR product |
|  | KO-RpHemY-5 | CGAAGCAGCTCCAGCCTACAGGTTACCGGCCGCCGCGC | *R. palustris* TIE-1 WT gDNA |
|  | KO-RpHemY-6 | AGGTCGACGGTATCGATAAGCTTGATATCGAATTCCTGCAGTTCTATTTGGCCCCCTGCCCCC | *R. palustris* TIE-1 WT gDNA |
| *R. sphaeroides* WT, Δ*mic60*, and Δ*orf52* verification | OUT2_F | CTACTTGACGCAGAGGCAGA | *R. sphaeroides* WT, Δ*mic60*, and Δ*orf52* gDNA |
|  | OUT2_R | GCTGGAGAGCCTTGCCTTTT | *R. sphaeroides* WT, Δ*mic60*, and Δ*orf52* gDNA |
|  | INT1_F | TTCGAAGCAGCTCCAGCCTAC | *R. sphaeroides* WT, Δ*mic60*, and Δ*orf52* gDNA |
|  | INT1_R | TGGAGAGCCTTGCCTTTTCC | *R. sphaeroides* WT, Δ*mic60*, and Δ*orf52* gDNA |
|  | INT2_F | TCGATGCCGAGAAAGGTGAG | *R. sphaeroides* WT, Δ*mic60*, and Δ*orf52* gDNA |
|  | INT2_R | TGTAGGCTGGAGCTGCTTCG | *R. sphaeroides* WT, Δ*mic60*, and Δ*orf52* gDNA |
|  | RSP6207_F | TGAGAAGAAACGGGACGGTG | *R. sphaeroides* WT, Δ*mic60*, and Δ*orf52* gDNA |
|  | RSP6207_R | ATTGCCTCAATGCGTTTGCG | *R. sphaeroides* WT, Δ*mic60*, and Δ*orf52* gDNA |
|  | RSP1508_F | GCTCTCCGCCAAGATGAAGT | *R. sphaeroides* WT, Δ*mic60*, and Δ*orf52* gDNA |
|  | RSP1508_R | GGCTCAACCGCTTTTTCGTC | *R. sphaeroides* WT, Δ*mic60*, and Δ*orf52* gDNA |
| *R. palustris* WT, Δ*mic60*, and Δ*orf52* verification | KOX_F1 | CGAAAAAGCGCTGCTCGAAG | *R. palustris* WT, Δ*mic60*, and Δ*orf52* gDNA |
|  | KOX_R1 | GCGCGATGATCACGAGAAAC | *R. palustris* WT, Δ*mic60*, and Δ*orf52* gDNA |
|  | KOX_R2 | GAACCTGCGTGCAATCCATC | *R. palustris* WT, Δ*mic60*, and Δ*orf52* gDNA |
|  | KOX_F3 | TCGCCTTCTTGACGAGTTCT | *R. palustris* WT, Δ*mic60*, and Δ*orf52* gDNA |
|  | KOX_R3 | CGAGCGACTTGGTGTCCTT | *R. palustris* WT, Δ*mic60*, and Δ*orf52* gDNA |
|  | KOY_F1 | GTCCCATCAATTTGCCACCG | *R. palustris* WT, Δ*mic60*, and Δ*orf52* gDNA |
|  | KOY_R1 | GCCGATGTGGTTTTGTCGAA | *R. palustris* WT, Δ*mic60*, and Δ*orf52* gDNA |
|  | KOY_R2 | GAACCTGCGTGCAATCCATC | *R. palustris* WT, Δ*mic60*, and Δ*orf52* gDNA |
| *R. sphaeroides mic60↑* | MitF-BamHI | GTGTGGATCCATGTCAGAACCGGAGTCCC | *R. sphaeroides* 2.4.1 WT gDNA |
|  | MitR-BglII | GTGTAGATCTCTTCCCTTCCGCGGCT | *R. sphaeroides* 2.4.1 WT gDNA |
| *R. sphaeroides orf52↑* | hemY_N-F-BamHI | GTGTGGATCCATGCTTTGGTCCTTGATCAAGATC | *R. sphaeroides* 2.4.1 WT gDNA |
|  | hemY_N-R-BglII | GTGTAGATCTTTTTGTGCTGGGCTCAACC | *R. sphaeroides* 2.4.1 WT gDNA |
| *R. palustris mic60↑* | pSRK-Mic60_F | ATAACAATTTCACACAGGAAACAGCATATGGTCGAGAACAGGCCCGAAC | *R. palustris* TIE-1 WT gDNA |
|  | pSRK-Mic60-6H_R | CCGGGTCGAATTTGCTTTCGAATTGCTAGCCTAGTGATGGTGATGGTGATGTGGCGACGGTTTTTGCA | *R. palustris* TIE-1 WT gDNA |
| *R. palustris orf52↑* | pSRK-HemY_F | ATAACAATTTCACACAGGAAACAGCATATGCTGCGCATCGTTCTGTTTCT | *R. palustris* TIE-1 WT gDNA |
|  | pSRK-HemY-6xH_R | CCGGGTCGAATTTGCTTTCGAATTGCTAGCTCAGTGATGGTGATGGTGATGGCGCGGCGGCCGGTAAC | *R. palustris* TIE-1 WT gDNA |
| *R. sphaeroides* overexpression verification by RT qPCR | qPCR_mic60_F | AACTCATGGCCGAAATGC | *R. sphaeroides* 2.4.1 cDNA from cells grown +/-IPTG |
|  | qPCR_mic60_R | CAAGAAGCAGCAGGACGAC | *R. sphaeroides* 2.4.1 cDNA from cells grown +/-IPTG |
|  | qPCR_orf52_F | GAGATCAACCCCAAGCACTC | *R. sphaeroides* 2.4.1 cDNA from cells grown +/-IPTG |
|  | qPCR_orf52_R | GACTTCATCTTGGCGGAGCG | *R. sphaeroides* 2.4.1 cDNA from cells grown +/-IPTG |
|  | qPCR_rpoz_F | ATGATCGAGAGCCACCAGAC | *R. sphaeroides* 2.4.1 cDNA from cells grown +/-IPTG |
|  | qPCR_rpoz_R | CACGCAGCAGCTTCTCTTC | *R. sphaeroides* 2.4.1 cDNA from cells grown +/-IPTG |
| *R. palustris* overexpression verification by RT qPCR | qPCR_mic60_F | GTTCCAGGCCTTGCTGAAG | *R. palustris* TIE-1 cDNA from cells grown +/-IPTG |
|  | qPCR_mic60_R | AGTTGGTCGATCAGCTCCTG | *R. palustris* TIE-1 cDNA from cells grown +/-IPTG |
|  | qPCR_orf52_F | GATCGACAAGAAGGCGTTTC | *R. palustris* TIE-1 cDNA from cells grown +/-IPTG |
|  | qPCR_orf52_R | CTTTCGTCGCTCTCTTCCAG | *R. palustris* TIE-1 cDNA from cells grown +/-IPTG |
|  | qPCR_rpoz_F | ACAAGAATCCCGTTGTTTCG | *R. palustris* TIE-1 cDNA from cells grown +/-IPTG |
|  | qPCR_rpoz_R | CGACCTCAACGAACTTCTGC | *R. palustris* TIE-1 cDNA from cells grown +/-IPTG |
